# Stage-resolved gene regulatory network analysis reveals developmental reprogramming and genes with robust stem-preferred expression in sorghum

**DOI:** 10.1101/2025.05.27.656329

**Authors:** Jie Fu, Brandon James, Madara Hetti-Arachchilage, Yingjie Lei, Brian McKinley, Evan Kurtz, Kerrie Barry, Stephen P. Moose, John E. Mullet, Kankshita Swaminathan, Amy Marshall-Colon

## Abstract

**Background:** Sorghum bicolor is a deep-rooted, heat- and drought-tolerant crop that thrives on marginal lands and is increasingly valued for its applications in biofuel, bioenergy, and biopolymer production. The sorghum stem, which can reach 4–5 meters in length, serves as the primary reservoir of both lignocellulosic biomass and soluble sugars, making it a promising bioenergy feedstock. Although recent advances in genetic, genomic, and transcriptomic resources have improved our understanding of sorghum biology, comprehensive genome-wide analyses of functional dynamics across diverse organ types and developmental stages remain limited. In particular, candidate genes with stem preferred expression pattern or their associated cis-regulatory elements, which may program key stem-related functions and enable organ- or tissue-specific engineering, have not yet been identified.

**Results:** To address these gaps, we reanalyzed a published RNA-seq dataset to identify genes with organ-preferential expression and to infer representative organ functions across major developmental stages. Our analysis revealed that the sorghum stem exhibits distinct temporal functional signatures, which correlate with the developmental dynamics of stem-specific genes and their associated regulatory elements. We further identified a set of genes with ubiquitous stem-specific expression across diverse sorghum genotypes, suggesting their universal importance and broad potential for genetic engineering applications. Among them, SbTALE03 and SbTALE04 emerged as stem hub transcription factors (TF). Both genes were empirically validated for their stem specificity across stages. Gene regulatory network analysis further indicated that these TFs participate in stage-specific transcriptional programs that maintain and regulate stem development.

**Conclusions:** This study presents a genome-wide analysis of organ-specific gene expression, functions, and regulatory networks in sorghum, with a focus on genes preferentially-expressed in stems and their promoter motifs. We identified a set of core stem-specific genes with ubiquitous expression across genotypes and developmental stages, including two experimentally validated transcription factors with potential roles in stem development. These findings offer valuable candidates for further functional characterization and genetic engineering aimed at improving sorghum stem biomass and composition.

## Background

*Sorghum bicolor* (L.) Moench is an important drought tolerant crop used to produce grain, feed, forage, and biofuels. Originally domesticated in Africa as a primary food source, sorghum was introduced to the United States in the 1750s [1,2]. Extensive breeding efforts during the 1900’s produced diverse genotypes and hybrids tailored for specific end uses. In the past two decades, biomass sorghum and sweet sorghum, characterized by high biomass yield and elevated stem sugar content, have garnered attention from both academic and industrial sectors for their potential in bioenergy and bioproduct production [3–9]. High biomass and sweet sorghum stems reach ∼4–5 m in length at maturity [10], account for approximately 80% of the plant’s total biomass [8], and store the majority of lignocellulosic biomass or soluble sugars [11,12].

Sorghum offers a compelling combination of traits that make it well suited for bioenergy production and sustainable agriculture. Like other C_4_ members of the *Poaceae* family, sorghum utilizes the C_4_ photosynthetic pathway [13,14], which confers high photosynthetic efficiency [3], drought and heat tolerance [15], and superior nitrogen-use efficiency [16], compared to C_3_ crops such as rice and wheat. Sorghum’s deep-rooting system [17,18] and low input requirements make the crop well suited for cultivation on marginal lands where water and nutrients are often limited [19,20]. Sorghum also holds several advantages over other C_4_ bioenergy crops such as maize, Miscanthus, and sugarcane [20,21]. Sorghum has a relatively small genome (∼730–800 Mbp) compared to other major grass crops, several reference quality genome sequences and extensive pan-genome resources [22–26], making it a valuable genetic/genomic model system for C_4_ grasses. Sorghum is typically cultivated as an annual crop but the species can also ratoon [27] by growing out tillers from the original stalks after harvest, providing the ecosystem services of a cover crop. These characteristics make sorghum an increasingly attractive crop for both research and sustainable production.

Plants are composed of distinct organs, each consisting of specialized tissues and cell types. While certain cell types are present in multiple organs, such as epidermal cells and vasculature containing phloem and xylem, each organ exhibits unique morphology and carries out specialized but complementary functions essential for plant growth and development [28]. Functional specializations are due in part to distinct gene expression programs that drive specific structural and physiological characteristics [29]. Thus, differential gene expression combined with translational and post-translational modifications can often explain cell, tissue, and organ functional specialization.

Next-generation sequencing (NGS) technologies make it possible to profile gene expression at the genome-wide level [30–34]. Although single-cell RNA sequencing and spatial transcriptomics have advanced our ability to resolve cell-level gene expression [35–40], bulk organ RNA sequencing remains a valuable source of information. Bulk RNA-seq enables a more accurate quantification of genes expressed at low levels which are frequently underrepresented or entirely missed in single-cell datasets due to gene dropout [41]. It also allows for robust comparisons between organs, making it an appropriate approach for investigating broader questions about organ-specific functions and development. For example, identifying genes that are specifically expressed in an organ across developmental stages provides insight into how organ identity is established, maintained, and regulated over time [29,42–44]. In addition, organ-specific genes also serve as a valuable source of information about promoter regulatory elements and transcription factor binding sites, which may be targets for sorghum trait improvement [45,46].

A large number of sorghum high-throughput datasets are publicly available [47–50]. Sreedasyam et al. [50] identified sorghum leaf-specific and seed-specific genes using the Tau metric, however, a much more comprehensive analysis of organ specific gene expression is needed. Other transcriptome studies have either focused on a single developmental stage [51], employed an inconsistent sampling strategy across multiple stages and organ/tissue types [49], concentrated on a specific gene family or biochemical pathway [52,53], or examined only a small set of genes [54–58]. Among these, Shakoor et al. [49] generated a valuable organ/tissue- and genotype-specific expression atlas using a microarray platform, but did not analyze regulatory mechanisms or provide genome-wide developmental trajectories of stem gene expression. As a result, important gaps remain in our understanding of gene expression dynamics throughout sorghum development and regulatory mechanisms underlying organ/tissue-specific expression.

Here, we utilized a published sorghum RNA-seq dataset encompassing major organ types, including stem, leaf, root, and seed, sampled across multiple developmental stages to identify organ-specific genes. We further assessed the consistency of stem-specific gene expression across a broader context using multiple sorghum genotypes. The expression of two candidate stem-specific genes with central roles in establishing and maintaining stem identity were experimentally validated. Our analysis of promoter-embedded regulatory elements uncovered stage-specific transcriptional dynamics. Finally, we constructed gene regulatory networks to investigate the functions and developmental dynamics of the two validated stem-specific regulators. This study advances our understanding of stem-specific gene regulation in sorghum and lays the groundwork for future development of targeted regulatory tools in bioenergy crop improvement.

## Methods

### RNA-seq datasets and data preprocessing

The RNA-seq dataset published by McCormick et al. [24] was used to identify organ-specific genes in sorghum, perform WGCNA-based analyses, and construct gene regulatory networks. The RNA-seq reads are available from the NCBI SRA under accessions SRA558272, SRA558514, and SRA558539. This dataset includes samples from various organ types (stem, leaf, root, panicle, peduncle, seed) collected at five developmental stages: the juvenile stage at 8 days after emergence (DAE), vegetative at 24 DAE, floral initiation at 44 DAE, anthesis at 65 DAE, and grain maturity at 96 DAE. All samples were collected from the grain sorghum genotype *Sorghum bicolor* BTx623. Developmental stages were defined based on morphological traits following Gerik et al. [59]. To focus the analysis on stem organ that is most relevant for bioenergy feedstock, panicle and peduncle samples, which are major reproductive stem-like organs, were excluded from analysis. Details of the 96 samples used in this study are provided in Supplemental Dataset 1. Genes with a mean transcripts per million (TPM) value ≥ 5 in at least one organ type were retained for downstream analyses.

To validate the stem specificity of genes identified in BTx623 across other genotypes, a secondary published RNA-seq dataset was used [60]. Gene expression profiles are available through the JGI genome portal under identifiers 1258721 and 1258409 (https://genome.jgi.doe.gov/portal/). This dataset includes leaf and stem samples collected from multiple sweet and bioenergy sorghum genotypes at three time points defined by days after planting (DAP): 57, 77, and 87 DAP, corresponding to vegetative, booting, and post-grain filling stage, respectively. Only genotypes with matched leaf and stem samples collected at the same developmental stage were included to assess organ-specific gene expression. The data structure and overview are shown in Figure 3a.

**Figure 1.**
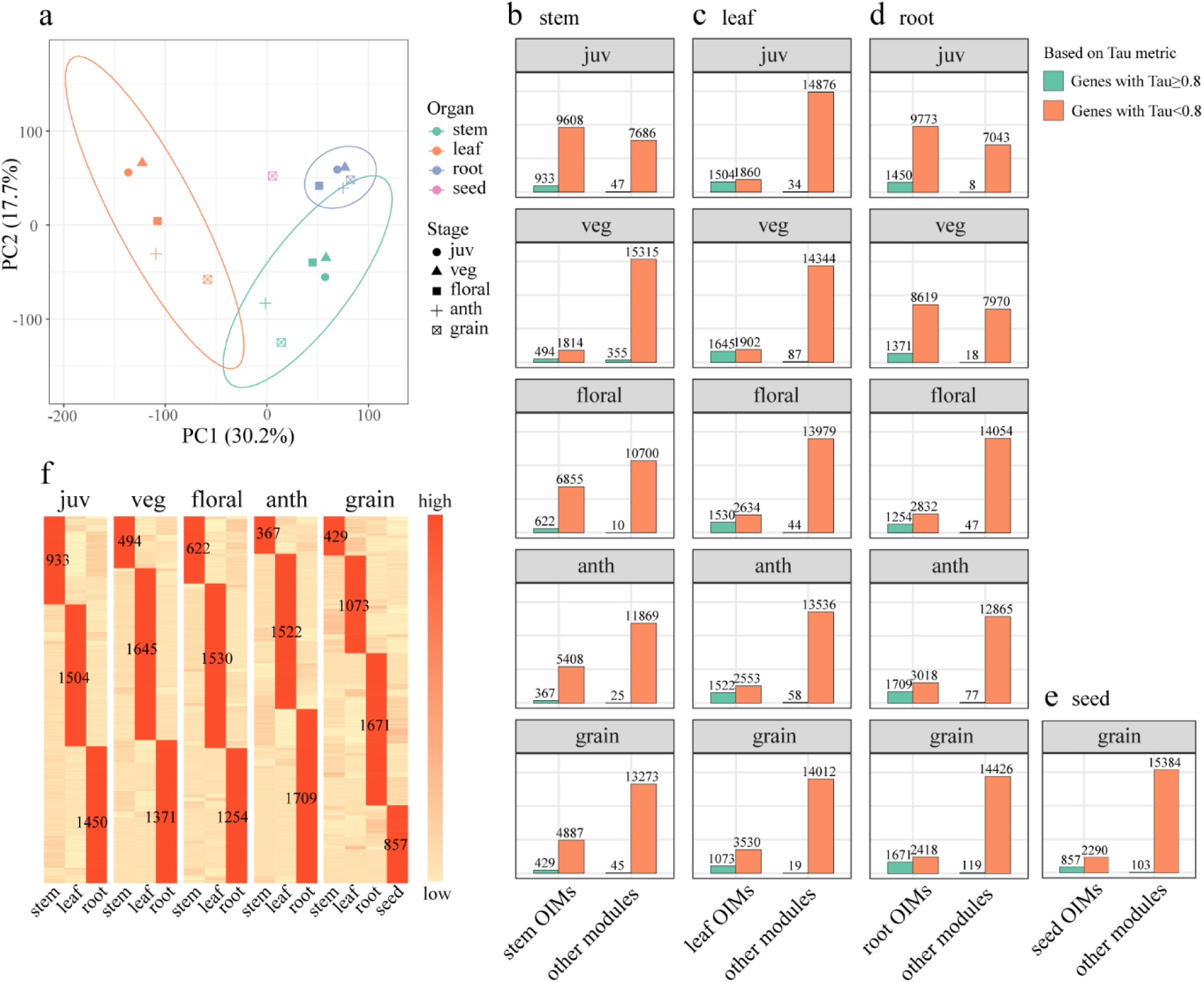
Identification of organ-specific genes in sorghum using integrated Tau metric and WGCNA. (a) Principal component analysis (PCA) of sorghum stem, leaf, root, and seed samples collected at five developmental stages: juvenile (juv), vegetative (veg), floral initiation (floral), anthesis (anth), and grain maturity (grain). (b–e) Distribution of genes with Tau-based organ-specific expression in organ-important modules (OIMs) versus other modules for: (b) stem, (c) leaf, (d) root, and (e) seed. (f) Heatmap of organ-specific gene expression. Mean expression (TPM) was centered and scaled by gene (rows).

**Figure 2.**
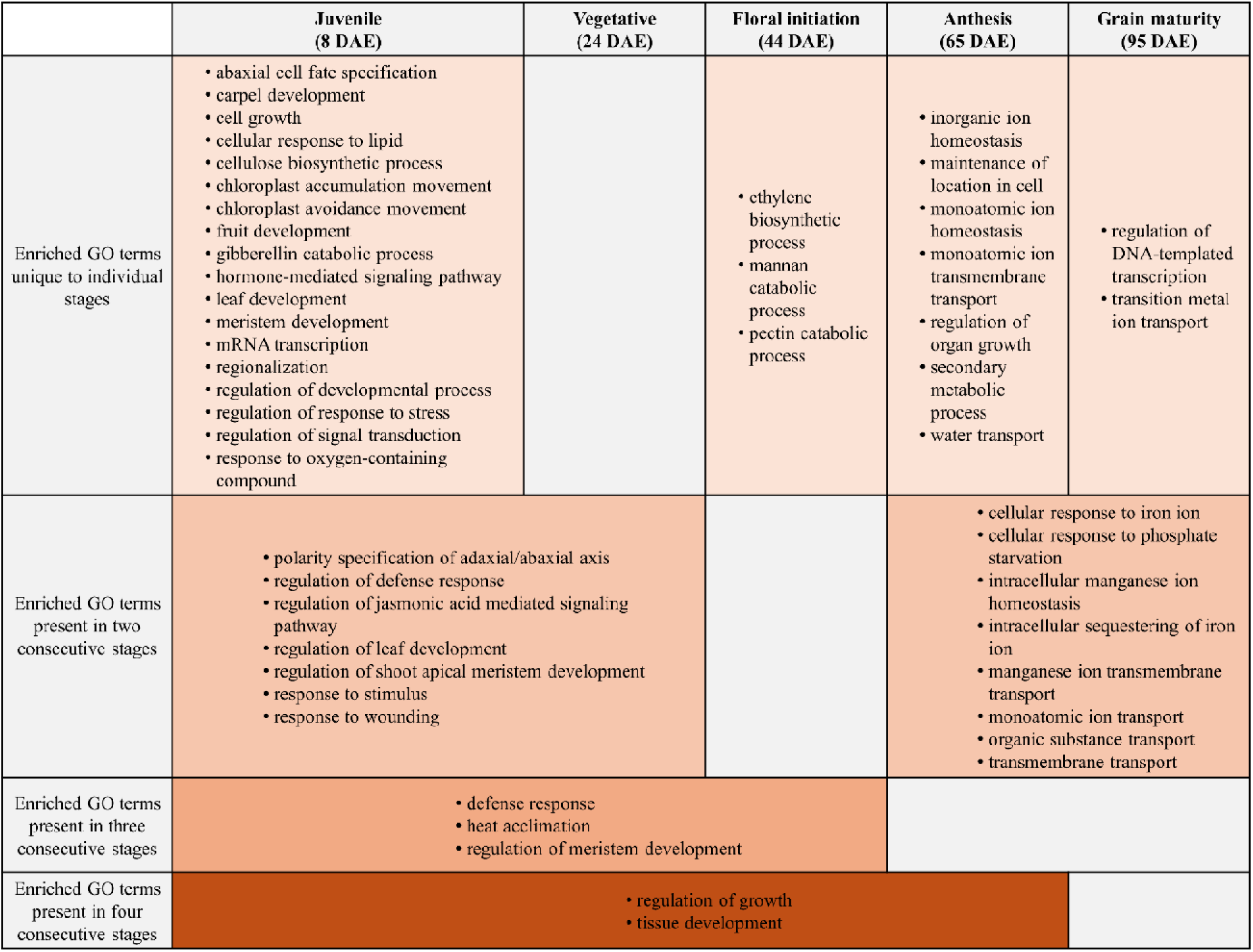
Unique and common enriched biological functions of stem-specific genes across developmental stages. Enriched Gene Ontology (GO) Biological Process (BP) terms presented include those that are unique to individual stages or found in at least two consecutive stages. GO terms listed here are only the most specific enriched terms (See Methods).

**Figure 3.**
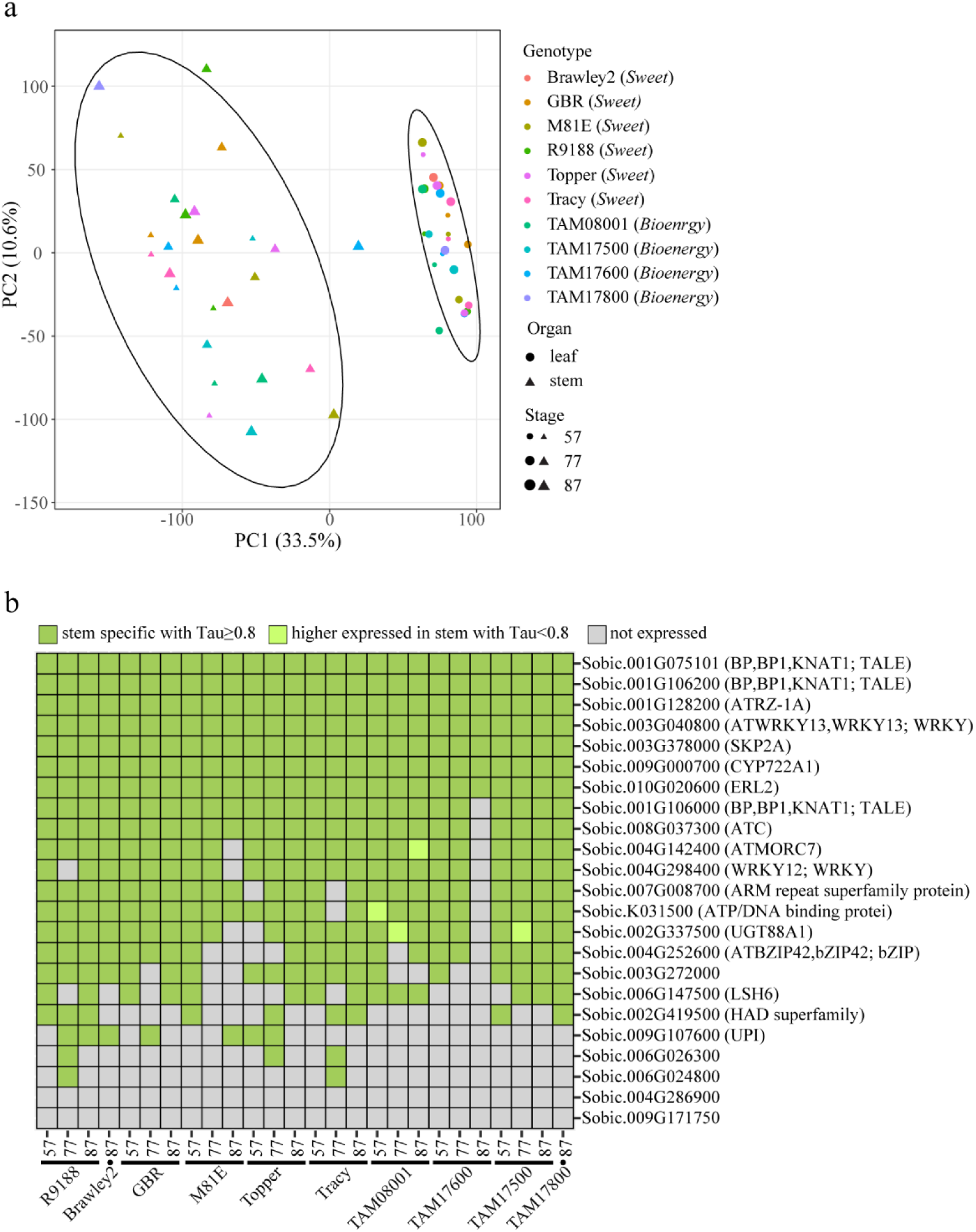
Stem specificity of genes in other sorghum genotypes across developmental stages. (a) Principal component analysis (PCA) of stem and leaf samples collected from multiple sorghum genotypes at 57, 77, and 87 days after planting (DAP). (b) Stem specificity across diverse sorghum genotypes for the ubiquitous stem-specific genes identified from BTx623. Organ specificity was estimated using the Tau metric.

### Tau metric to evaluate gene organ specificity

The Tau index has been demonstrated as a robust method for evaluating gene expression specificity in sorghum [37], as well as other organisms and circumstances [61,62]. We applied the Tau metric to the two published RNA-seq datasets mentioned above to identify and validate genes preferentially expressed in sorghum stem organ. Tau value was calculated using the organs collected from the same sorghum genotype at a certain growing stage, thus the resulting Tau value is genotype-, stage-, and dataset-specific (e.g., stem-specific genes for BTx623 at vegetative stage based on RNA-seq data from McCormick et al. [24]). The Tau calculation method and gene specificity criteria are described and validated in Kryuchkova-Mostacci & Robinson-Rechavi [61]. Briefly, each gene was assigned a Tau value between 0 and 1, with higher values indicating stronger organ specificity and lower values indicating ubiquitous expression. Genes with Tau values between 0.8 and 1.0 were classified as preferentially expressed (0.8 ≤ Tau < 1) or exclusively expressed (Tau = 1), with specificity assigned to the organ type showing the highest expression.

### WGCNA co-expression network construction and module detection

Co-expression networks were constructed using the WGCNA package (1.72-5) in R environment (4.3.3). Preprocessed RNA-seq data for sorghum BTx623, including individual replicates, were log2-transformed to meet the normality assumption, verified by the Shapiro– Wilk test. A signed co-expression network was created for each developmental stage by (1) using gene pair Pearson correlation coefficient (PCC) for adjacency matrix calculation by raising |(1 + PCC)/2| to a soft threshold power and (2) calculating the topological overlap matrix (TOM) from the adjacency matrix. Soft thresholds were set to 14 for juvenile, 16 for vegetative, 13 for floral initiation, 14 for anthesis, and 16 for grain maturity stage to meet the scale-free network assumption. To identify co-expression modules among genes with similar expression profiles, we applied hierarchical clustering using distance matrix from TOM (i.e., 1–TOM). We here aimed for smaller module sizes, as they are more effective for identifying gene groups associated with specific biological functions [63]. Parameters controlling module size were set as follows: *minClusterSize = 15, deepSplit = 4, and cutHeight = 0.1*. Detailed module information and gene module membership (MM) are provided in Supplemental Dataset 9.

### Identification of organ-important modules

We defined “organ-important modules” as WGCNA co-expression modules that are enriched for genes with organ-preferential expression. Module importance for a given organ type was estimated using point-biserial correlation (r b) between module eigengene expression, a continuous variable, and organ identity, a binary variable. For example, to estimate module importance for stem organ, stem samples were coded as 1 and all other samples as 0. Modules showing a significantly positive correlation (r b > 0 & *p* < 0.05) between eigengene expression and stem identity were considered stem-important. These organ-important modules consist of genes that are highly expressed in a given organ type, as indicated by high eigengene values in stem samples (Figure S9), but also exhibit a shared expression pattern across organ types, a fundamental property of WGCNA modules. In terms of identifying organ-specific genes, this network-based approach incorporates replicate variation and co-expression structure, complementing the Tau metric, which is based solely on mean expression values. For genes with high Tau values (≥ 0.8), those in organ-important modules tend to have higher Tau values than those in non-organ-important modules (Figure S10), supporting the coherence and reliability of both approaches. Module importance for each organ type is provided in Supplemental Dataset 10.

### Stem hub transcription factor (TF) identification

Stem hub TFs were identified based on: (i) classification in the Plant Transcription Factor Database (https://planttfdb.gao-lab.org/) as a sorghum transcription factor; (ii) presence in stem-important modules defined previously; and (iii) high gene significance (GS) relative to stem identity (GS > 0.8 & *p* < 0.05). GS represents the correlation between gene expression and binary stem identity across samples. We did not apply gene membership (MM) within stem-important modules or intramodular connectivity (KIM), as prior work shows GS correlates well with both parameters [64] and our goal here is to recover a broader set of potential stem hub TFs. The identified stem hub TFs are listed in Supplemental Dataset 7.

## GO enrichment and identification of representative GO terms

GO enrichment analysis was conducted using the gprofiler2 package (0.2.2) in R environment (4.3.3). To utilize the package’s built-in gene annotation for *Sorghum bicolor*, gene identifiers with the “*Sobic.*” prefix were converted to the Ensembl format “*SORBI_3*,” which is recognized by gprofiler2. Annotations supported exclusively by electronic evidence (IEA) were excluded to enhance biological reliability. The background gene set consisted of all expressed genes, excluding lowly expressed ones filtered during data preprocessing. Multiple testing correction was performed using the false discovery rate (FDR), and GO terms with FDR < 0.05 were considered significantly enriched. To reduce redundancy among enriched terms, a hierarchical GO term network was constructed by linking each term to its direct parent and child nodes. The most specific child terms, those located at the lowest level of the hierarchy, were selected to represent distinct functional groups and were used to generate visualizations, including Figure 2 and S2.

### Phylogenetic analysis

*Sorghum bicolor* genes that encode TALE transcription factors (SbTALEs), including KNOX and BEL1-like transcription factors, were previously identified and numbered based on a phylogenetic analysis [65]. In this study, protein sequences of SbTALEs that encode KNOX family transcription factors [65] and KNOX family (KNOX clades I-III) protein sequences from Arabidopsis thaliana [65–68], and Zea mays [66,67] were collected from *A. thaliana* TAIR10, *O. sativa* v7.0, *S. bicolor* v3.1.1. and Z*. mays* v5.0 genome sequences in Phytozome v13. KNOX family protein sequences from rice, maize and Arabidopsis were used to identify sorghum homologs using BLAST analysis. In addition to the 23 SbTALE genes identified previously [65], two additional sorghum genes, Sobic.002G314000 and Sobic.003G331500, were identified as best hits via BLAST homology searches on Phytozome v13 to Zm00001eb322410 (ZmTALE32) and Zm00001eb013560 (ZmTALE4) respectively (E-values = 9 x 10^-32^ and 2.5 x 10^-31^ respectively). KNOX family sequences were aligned using the multiple sequence alignment MUSCLE v5.1.0 algorithm. Model selection was conducted using ModelTest-NG v0.1.7, with the best protein substitution model determined by the lowest corrected Akaike Information Criterion (AICc) value. Phylogenetic analyses were performed using RAxML v8.2.12 under the JTT protein substitution model, with a gamma distribution to account for rate variation.

Maximum likelihood (ML) analysis included 1,000 bootstrap replicates to assess branch support. Phylogenetics, alignment, and model selection were performed on the Grace supercomputer at Texas A&M High-Performance Research Computing (HPRC). The resulting trees were visualized and annotated using Dendroscope3.

### Motif enrichment analysis, positional preference and similarity estimation

Known motifs from the JASPAR 2020 Core Plant Non-Redundant database were scanned within 1.5 kb promoter regions upstream of the ATG start codon in organ-specific genes using Find Individual Motif Occurrences (FIMO) in MEME suite (*p* < 1e-4). A motif was considered enriched if genes with the motif were significantly overrepresented in organ-specific genes compared to non-specific genes, as determined by Fisher’s exact test (*p* < 0.05). To examine the positional preference of enriched known motifs along the promoter region, promoter sequences of organ-specific genes were shuffled using the “shuffle_sequences” function from the universalmotif package (default settings), divided into 100 bp bins, and scanned for motif occurrences using FIMO (*p* < 1e-4). A higher ratio of observed motif occurrences in the original promoters to expected occurrences in the shuffled sequences indicates stronger positional preference. *de novo* enriched motif discovery was performed using MEME (5.5.1), with motif lengths ranging from 5 to 21 bp and an *E-value* threshold of < 0.05. Background Markov models were generated using RSAT (https://rsat.eead.csic.es/plants/create-background-model_form.cgi). All analyses were conducted under the Unix environment. To assess whether *de novo* enriched motifs were represented in the known motif database, similarity between *de novo* and known motifs was evaluated by calculating Pearson correlation coefficients (PCC) using universalmotif package (1.20.0) in R (4.3.3). *de* novo enriched motifs with a maximum PCC of 0.49 or lower against any known motif, which corresponds to the 95th percentile of PCC values within the known motif database, were considered novel motifs not currently represented in the existing database. Their MEME format representations are provided in Supplemental Dataset 5. This PCC threshold is based on the assumption that most motifs in the database are nonredundant and exhibit low similarity to each other.

### Gene regulatory network (GRN) construction

GRN was constructed by integrating transcription factor (TF)–TF binding site (TFBS) regulatory interactions with gene co-expression relationships. TF–TFBS regulatory interactions were predicted using the web-based PlantPAN4.0 “Promoter Analysis” tool, which scanned 1.5 kb promoter regions upstream of the ATG start codon for TFBS. TFs from specific families were assumed to regulate genes containing the corresponding TFBS in their promoters, serving as the first layer of biologically relevant evidence for GRN construction. These predicted interactions were stored as a directed edge table, with TFs as regulators (i.e., source nodes) and genes containing the corresponding TFBS within their promoters as targets (i.e., target nodes). Co-expression relationships, serving as the second layer of information derived from expression data, were assessed using Pearson correlation coefficients (PCC) based on stem-only expression data at each developmental stage, excluding other organ types. Other organ types were excluded to preserve subtle stem-specific correlations that might be obscured across organs. Gene pairs with PCC > 0.8 and *p* < 0.05 were included in an undirected edge table. The final GRN was generated by retaining edges that were present in both the directed and undirected edge tables.

To construct specific networks for the two stem hub TFs focused by this study, Sobic.001G106000 (SbTALE03) and Sobic.001G106200 (SbTALE04), the full GRN was further filtered to include only: (1) regulatory interactions where either of the two TFs served as the source node, and (2) target genes that were located within the same WGCNA co-expression module as the respective TF, thereby capturing their highly co-expressed and functionally associated targets. The resulting SbTALE03 and SbTALE04 networks are illustrated in Figure 8b, and the corresponding node and edge data are provided in Supplemental Dataset 8.

### RNAscope *in situ* hybridization (ISH) visualization and quantification

Stem organs were collected from *Sorghum bicolor* cv. BTx623 at vegetative and post-floral initiation stages for RNAscope ISH. Sample fixation, embedding, and pretreatment followed the RNAscope 2.5 FFPE Sample Preparation and Pretreatment protocol (https://acdbio.com/sites/default/files/UM%20322452%20FFPE%20SamplePrep%20Rev%2003152017_0.pdf), with the following modifications: 10 μm paraffin sections were cut and baked for 2 h; slides were dried at 60 °C for 20 min after deparaffinization. Gene-specific probes were hybridized using the RNAscope Multiplex Fluorescent Reagent Kit v2 (Cat# 323100) and visualized with Opal 520 dye and DAPI counterstain. Imaging was performed using a Nikon AXR confocal microscope with a resonant scanner at 20× magnification. Additional protocol details are available in Fu et al. [37]. Transcript counts were obtained using the Nikon Elements Software analysis tool Object Count with the following settings: (Leaves) Threshold Multichannel 1,300 to 15,000; EqDiamter 0.50 to 3.00; Circularity 0.80 to 1.00; (Stems) Threshold Multichannel: 650 to 9,000; EqDiamter 0.50 to 3.00; Circularity 0.80 to 1.00; (Roots) Threshold Multichannel: 1,500 to 10,000; EqDiamter 0.50 to 3.00; Circularity 0.80 to 1.00. To account for differences in background expression levels across organs, gene counts were normalized using the bacterial *dapB* gene as the negative control, which was assumed to show no expression after normalization.

### Protein purification

Coding sequence was codon-optimized and synthesized by GenScript, and subsequently cloned into the pET-28a(+) expression vector at the BamHI and EcoRI sites. The resulting construct was transformed into *E. coli* BL21(DE3) cells. A 200 mL bacterial culture was grown in LB medium and induced with 0.1 mM IPTG at 20 °C with shaking at 210 rpm for 24 h. Cells were harvested by centrifugation at 8000 × g for 10 min at 4 °C, and the pellet was resuspended in 20 mL lysis buffer containing: 50 mM phosphate buffer (pH 8.0), 300 mM NaCl, 10% glycerol, 1% Triton X-100, 25 mM imidazole, 5 mM MgCl , 50 U/mL Pierce™ Universal Nuclease for Cell Lysis (Thermo Fisher, 88700), and 1× Halt™ Protease Inhibitor Cocktail (Thermo Fisher, 87785). Cell lysis was performed by ultrasonication on ice. The lysate was clarified by centrifugation, and the supernatant was incubated with HisPur™ Ni-NTA Spin Columns (Thermo Fisher, 88226) at 4 °C for 30 min. The columns were washed with 50 mL wash buffer (50 mM phosphate buffer pH 8.0, 300 mM NaCl, 10% glycerol, 0.5% Triton X-100, 75 mM imidazole, 5 mM MgCl ), and the bound protein was eluted with 3 mL elution buffer containing 50 mM phosphate buffer (pH 8.0), 300 mM NaCl, 10% glycerol, 0.5% Triton X-100, 300 mM imidazole, and 5 mM MgCl . Protein concentration was determined by measuring absorbance at 280 nm, and purity was assessed by SDS-PAGE.

### Electrophoretic Mobility Shift Assay (EMSA)

Promoter sequence was defined as 1.5 kb upstream of ATG start codon and obtained from Phytozome using Biomart tool. Sequences were amplified using either a 5′ IRD700-labeled forward primer (for the labeled probe) or an unlabeled forward primer, along with an unlabeled reverse primer. PCR amplification was carried out using PrimeSTAR® Max DNA Polymerase Ver.2 (Takara, R047A). The PCR products were run on a 1% agarose gel and purified using the GeneJET Gel Extraction Kit (Thermo Fisher, K0691). For the binding assay, 50 ng of the labeled probe or 5 μg of the unlabeled probe was incubated with 10 μg of purified SbTALE03 protein in a 20 μL reaction containing 1× binding buffer, 5% glycerol, 5 mM MgCl , 0.5% Triton X-100, and 1μL poly(dI-dC) (Thermo Fisher, 20148X). The mixture was incubated at room temperature for 20 minutes. Following incubation, the samples were mixed with loading buffer and subjected to electrophoresis on a 5% TBE polyacrylamide gel (Bio-Rad, 4565014) at 100 V for 50 minutes. Bands were visualized by direct scanning using the Odyssey CLx Imaging System.

## Results

### Identification of sorghum organ-specific genes

RNA-seq data enables discovery of genes with distinctive expression patterns. In this study, we used an RNA-seq dataset from McCormick et al. [24], which includes profiles of samples from different sorghum organs (leaf, stem, root, seed) collected at five stages of development (juvenile, vegetative, floral initiation, anthesis, grain maturity) (Figure 1a and Supplemental Dataset 1). The data was used to identify genes preferentially expressed in specific organ types at each developmental stage by integrating the Tau index [61,62] and Weighted Gene Co-expression Network Analysis (WGCNA). The Tau-based analysis revealed that 21–23% of expressed genes exhibited high organ specificity (Tau ≥ 0.8) at each stage of plant development (Figure S1). Notably, a significant majority (93 ± 10%; Fisher’s exact test, *p* < 0.001) of these genes were located within WGCNA network modules that were highly correlated with specific organ types (Figure 1b–e). These modules were defined by a significant and positive point-biserial correlation (r_p_b) between the module eigengene and organ identity (r_p_b > 0 and *p* < 0.05); such modules are hereafter referred to as “organ-important modules”. The coherence between Tau values and WGCNA-derived organ-important modules indicates that Tau analysis provides a reliable way to identify genes with organ specific expression in sorghum. Genes meeting both criteria, Tau ≥ 0.8 and inclusion in organ-important modules, were classified as organ-specific genes (Figure 1f and Supplemental Dataset 2).

Organ identity and function are correlated with unique patterns of gene expression, biochemical pathways, and other molecular signatures. Enriched Gene Ontology (GO) terms of organ-specific genes revealed that these genes encode distinct functions in stems, roots, leaves and seeds (Figure S2 and Supplemental Dataset 3). For example, the stem is primarily associated with the GO terms of defense and transport; the leaf with photosynthesis-related processes; the root with cell wall organization, sulfation, and glutathione metabolism; and the seed with protein oligomerization, stress response, and lipid storage (Figure S2). Of these organs, the least is known about sorghum stem molecular function and regulation.

### Dynamic expression of stem-preferred genes during plant development

The analysis of stem-preferred expression revealed that specific biological processes are temporally enriched in the stem at different stages of plant development (Figure 2). During the juvenile and adult vegetative stage, stem-specific genes are enriched for GO terms associated with developmental regulation and stress responses, including polarity specification, shoot apical meristem (SAM) development, leaf development, and defense signaling pathways such as jasmonic acid response and wounding (Figure 2). These functions indicate active tissue patterning and protective mechanisms when the stem is growing and structurally dependent on the surrounding leaf sheath for mechanical support.

At floral initiation, the transition from vegetative to reproductive growth, stem-preferred gene expression indicates there is an increase in ethylene biosynthetic processes as well as mannan and pectin catabolic processes (Figure 2). Concurrent enrichment in cell wall remodeling pathways may support rapid stem elongation during the “booting” phase.

At anthesis and grain maturity when stem growth is no longer occurring, stem-specific genes are involved in nutrient transport and homeostasis, including responses to iron and phosphate, intracellular ion regulation, and transmembrane transport (Figure 2). These results may reflect the stem’s role in long-distance transport and nutrient reallocation, as photosynthates and solutes from the leaves, stems, and roots are mobilized and transported through stems to support grain development after anthesis.

The changes in stem-preferred gene expression during development underscore the dynamic nature of stem function.

### Stem-specific gene expression in diverse sorghum genotypes

Genes with consistent stem specific expression across genotypes and stages of development could have broad applications in sorghum stem engineering. Twenty-three genes were identified that exhibit stem-specific expression across all developmental stages (Figure S3 and Supplemental Dataset 4). These genes may play essential roles in maintaining stem identity and supporting fundamental, stage-independent stem functions.

Among these potential ubiquitous stem-specific genes, Sobic.001G106000, Sobic.001G106200, and Sobic.001G075101 belong to the Three Amino acid Loop Extension (TALE) homeodomain transcription factor family and are orthologous to KNOTTED-LIKE FROM ARABIDOPSIS THALIANA 1 (KNAT1, AT4G08150) (Supplemental Dataset 4), which is known to regulate meristematic cell identity. Other transcription factors with ubiquitous stem-specific expression include members of the WRKY family (Sobic.003G040800, Sobic.004G298400) and the bZIP family (Sobic.004G252600) (Supplemental Dataset 4), suggesting broad regulatory functions across stem tissues. In addition to these TFs, Sobic.002G337500, homologous to AT3G16520 (Supplemental Dataset 4), encodes UDP-glucosyl transferase 88A1, which may function in cell wall modification; Sobic.004G142400, homologous to AT4G24970 (Supplemental Dataset 4), encodes a histidine kinase potentially involved in signal transduction.

The expression of genes with stem specific expression in the grain sorghum genotype BTx623 was evaluated in a diverse panel of sweet and bioenergy sorghum genotypes. We applied Tau analysis on an RNA-seq dataset that includes stem and leaf samples collected from different sorghum genotypes during three stages of plant development, vegetative phase at 57 days after planting (DAP), during the booting phase (77 DAP), and post-grain filling (87 DAP) (Figure 3a). About 70% (16/23) of the ubiquitous stem-specific genes identified in BTx623 over development showed stem specific expression in ∼73% (19/26) of genotype-stage combinations included in this multi-genotype dataset (Figure 3b). Furthermore, seven genes consistently exhibited stem specific expression across all genotypes and stages of development (Figure 3b and Supplemental Dataset 4).

### Characterization of stem specific expression of SbTALE03 and SbTALE04

Two genes (Sobic.001G106000, Sobic.001G106200) that showed stem organ specific expression were characterized in greater depth. Both genes encode homeodomain TALE transcription factors. Phylogenetic analysis of sorghum, rice, and maize TALE transcription factors showed that Sobic.001G10600 (*SbTALE03*) was most closely related to *ZmTALE12* (Zm00001eb055940) and Sobic.001G0106200 (*SbTALE04*) was most closely related to *ZmTALE11* (Zm00001eb055920) (Figure S4).

Besides showing stem specificity in BTx623 (Figure 4a), *SbTALE03* and *SbTALE04* were expressed in apical tissue of growing stems, and the stem nodal plexus, internode, stem intercalary meristem (IM), and pulvinus tissues of stem node-internode segments collected at a stage where internodes were >50% elongated in biomass sorghum TX08001 (Figure 4b). The genes were also expressed in the stem peduncle that is produced between floral initiation and anthesis (Figure 4a). Expression of the genes was not detected in leaf blade, leaf sheath or root tissues (Figure 4a and 4b).

**Figure 4.**
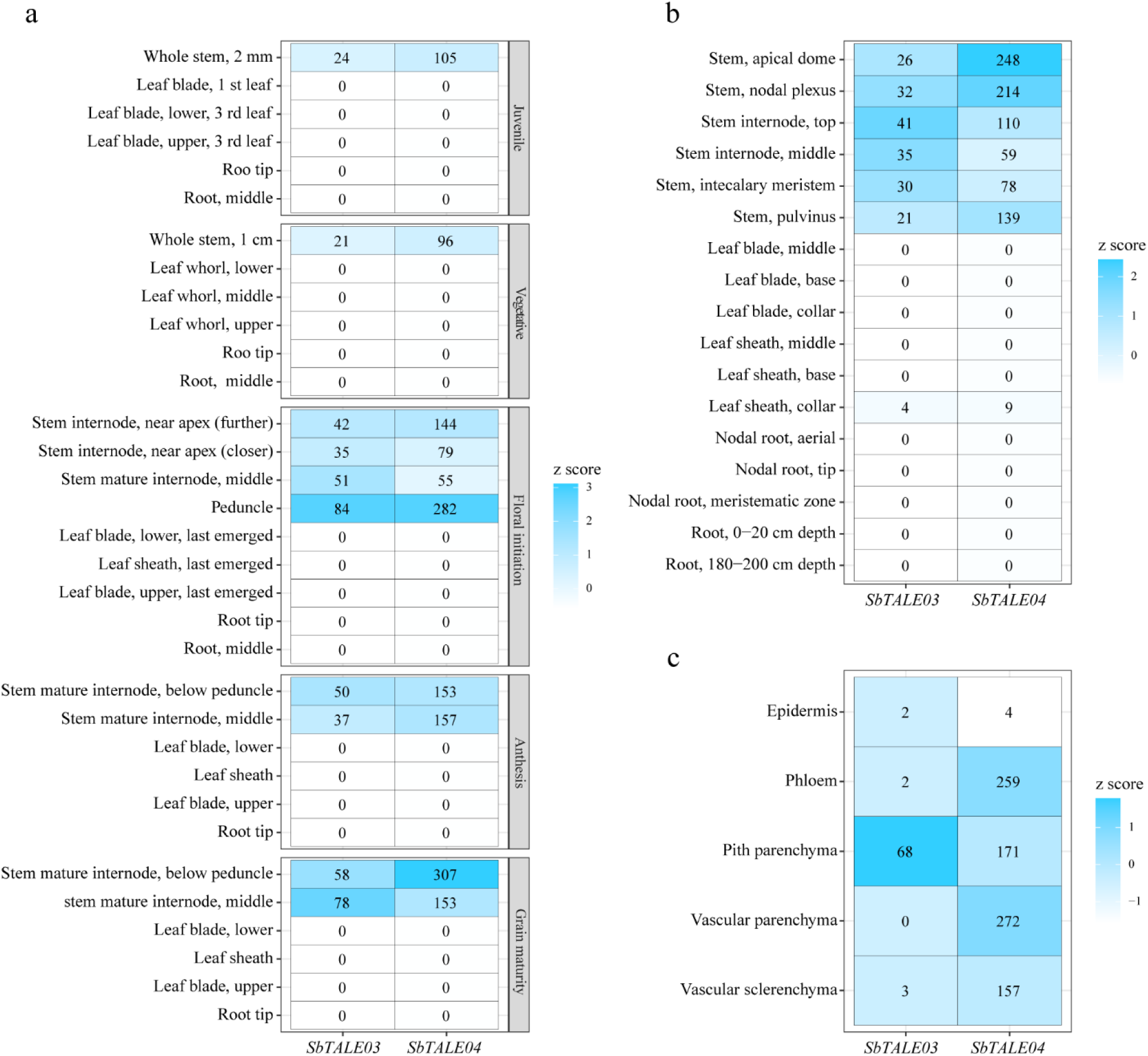
Transcript expression heatmaps of *SbTALE03* and *SbTALE04*. (a) Gene expression profiles across developmental stages in grain sorghum BTx623. (b) Expression profiles in biomass sorghum TX08001 at the vegetative stage. (c) Stem cell-type-specific expression in sweet sorghum Wray at the vegetative stage. Detailed sample description for panel a is in Supplemental Dataset 1.

Expression of *SbTALE03* and *SbTALE04* in stem cell types was characterized using previously published transcriptome profiles of vegetative phase stem internode cell types [37]. The analysis showed that *SbTALE03* was differentially expressed in pith parenchyma cells whereas *SbTALE04* was expressed in stem pith parenchyma, phloem, vascular parenchyma and vascular sclerenchyma/fiber cells (Figure 4c).

To experimentally validate stem-specific expression patterns identified by RNA-seq, we performed RNAscope *in situ* hybridization (ISH) in sorghum across developmental stages for *SbTALE03* and *SbTALE04*. RNAscope results revealed markedly higher transcript abundance of both genes in BTx623 stem organs compared to leaf and root at both vegetative (Figure 5) and post–floral initiation stages (Figure 6). In non-stem organs, expression was either undetectable or observed at 5.7-to 26.6-fold lower levels than in stem (Figure 5e and 6e). Beyond organ specificity, the relative expression levels and developmental dynamics of the two genes were consistent between RNA-seq and RNAScope results. Specifically, *SbTALE04* showed higher expression than *SbTALE03* at both tested stages (Figure 4a, 5e, and 6e). While *SbTALE04* expression remained stable over time, *SbTALE03* expression increased (Figure 4a, 5e, and 6e). In addition, in RNAscope images, at vegetative stage, *SbTALE03* only shows signals in stem pith parenchyma cells (Figure 5a), while *SbTALE04* shows signals in both pith parenchyma and vascular bundle tissues such as phloem cells (Figure 5b). This stem cell-type-specific expression pattern is consistent with stem cell type transcriptome analysis of vegetative-phase Wray (sweet sorghum) and Tx430 (grain sorghum) fully elongated stem internodes reported by Fu et al. [37].

**Figure 5.**
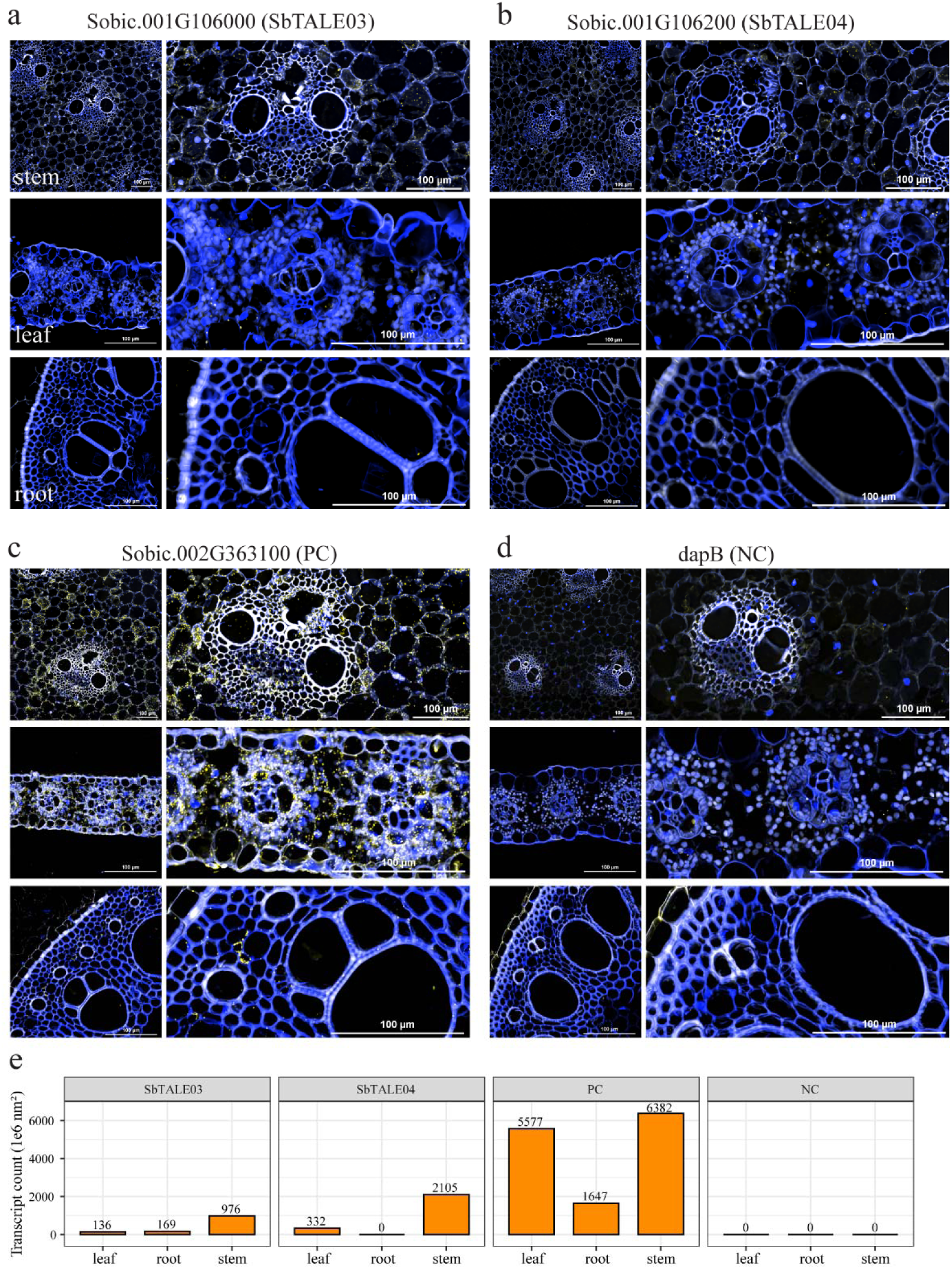
RNAscope ISH validation of *SbTALE03* and *SbTALE04* at the vegetative stage in sorghum BTx623. (a) *SbTALE03*; (b) *SbTALE04*; (c) Positive control (PC) probed for *Sobic.002G363100; SbEIF4*; (d) Negative control (NC) probed for *dapB*. (e) Transcript quantification. Transcript signals appear as small yellow puncta.

**Figure 6.**
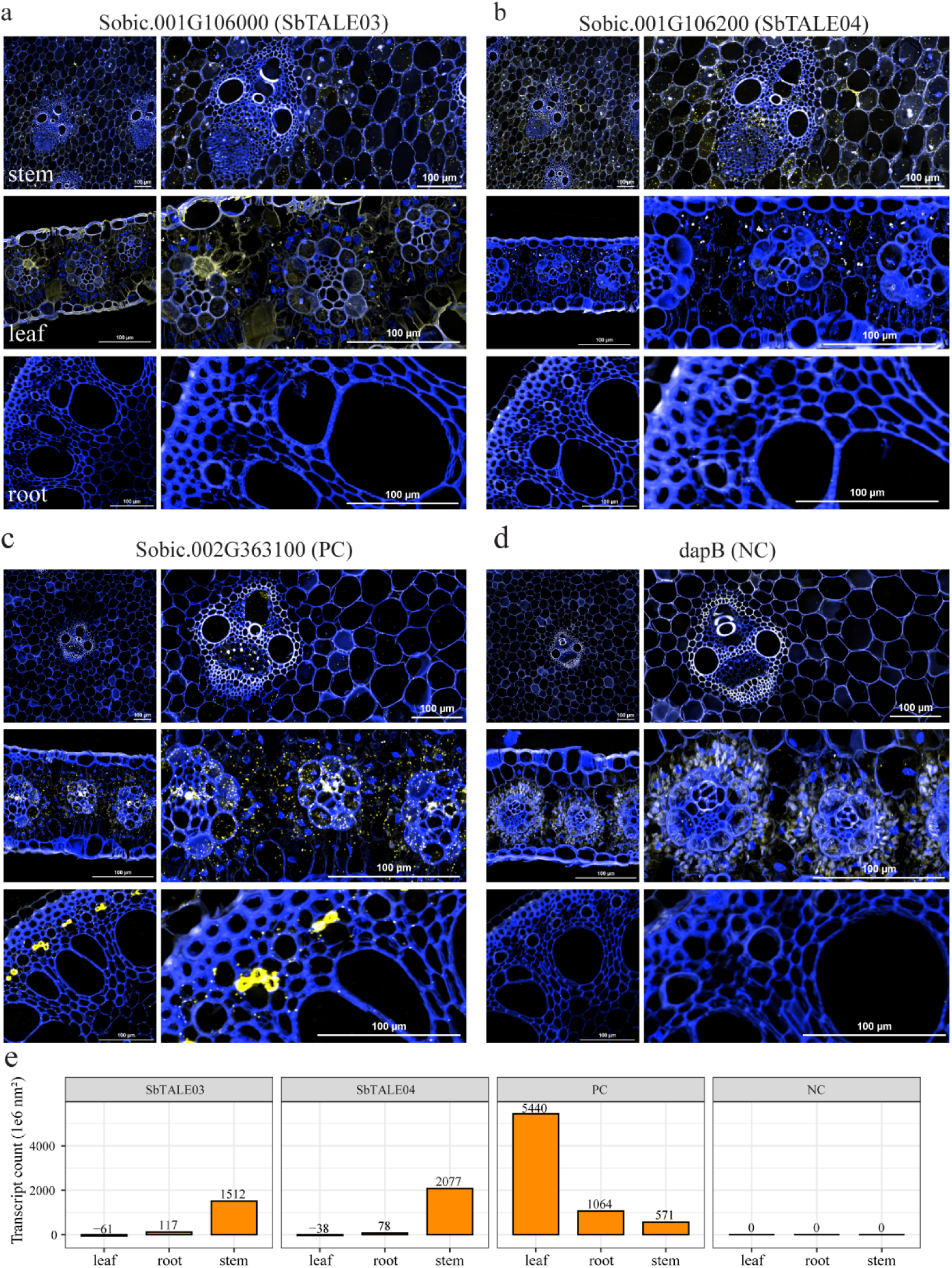
RNAscope ISH validation of *SbTALE03* and *SbTALE04* at the post-floral initiation (post-FI) stage in sorghum BTx623. (A) *SbTALE03*; (B) *SbTALE04*; (C) Positive control (PC) probed for *Sobic.002G363100; SbEIF4*; (D) Negative control (NC) probed for *dapB*. (e) Transcript quantification. Transcript signals appear as small yellow puncta.

The RNA-seq data from diverse sorghum genotypes and the consistency between RNA-seq and RNAscope results support the robustness of our methodology for identifying stem-specific genes. *SbTALE03* and *SbTALE04*, along with other high-confidence, genes with stem-specific expression (Figure 3b and Supplemental Dataset 4), represent promising candidates for functional analysis and targeted stem engineering tools in sorghum.

### Motif analysis of organ-specific gene promoters

Transcription factor binding sites (TFBS) in promoter regions are sites where transcription factors bind and mediate organ-specific transcriptional programs. To investigate potential regulatory motifs underlying organ-specific gene expression, we first searched for known motifs from available databases within the 1.5 kb region upstream of the ATG start codon of organ-specific genes. For each motif, we applied Fisher’s exact test to determine whether genes containing that motif were significantly enriched in a specific organ-specific gene set compared to all other expressed genes (*p* < 0.05). This test is based on the hypothesis that if regulatory motifs contribute to organ-specific gene expression, then genes with similar expression patterns should share common upstream motifs more frequently than expected by chance [69]. Our results indicate that these enriched known motifs are most frequently located within ∼100–300 bp of the start codon (Figure 7a), suggesting this region may play a particularly important role in organ-specific regulation in sorghum.

**Figure 7.**
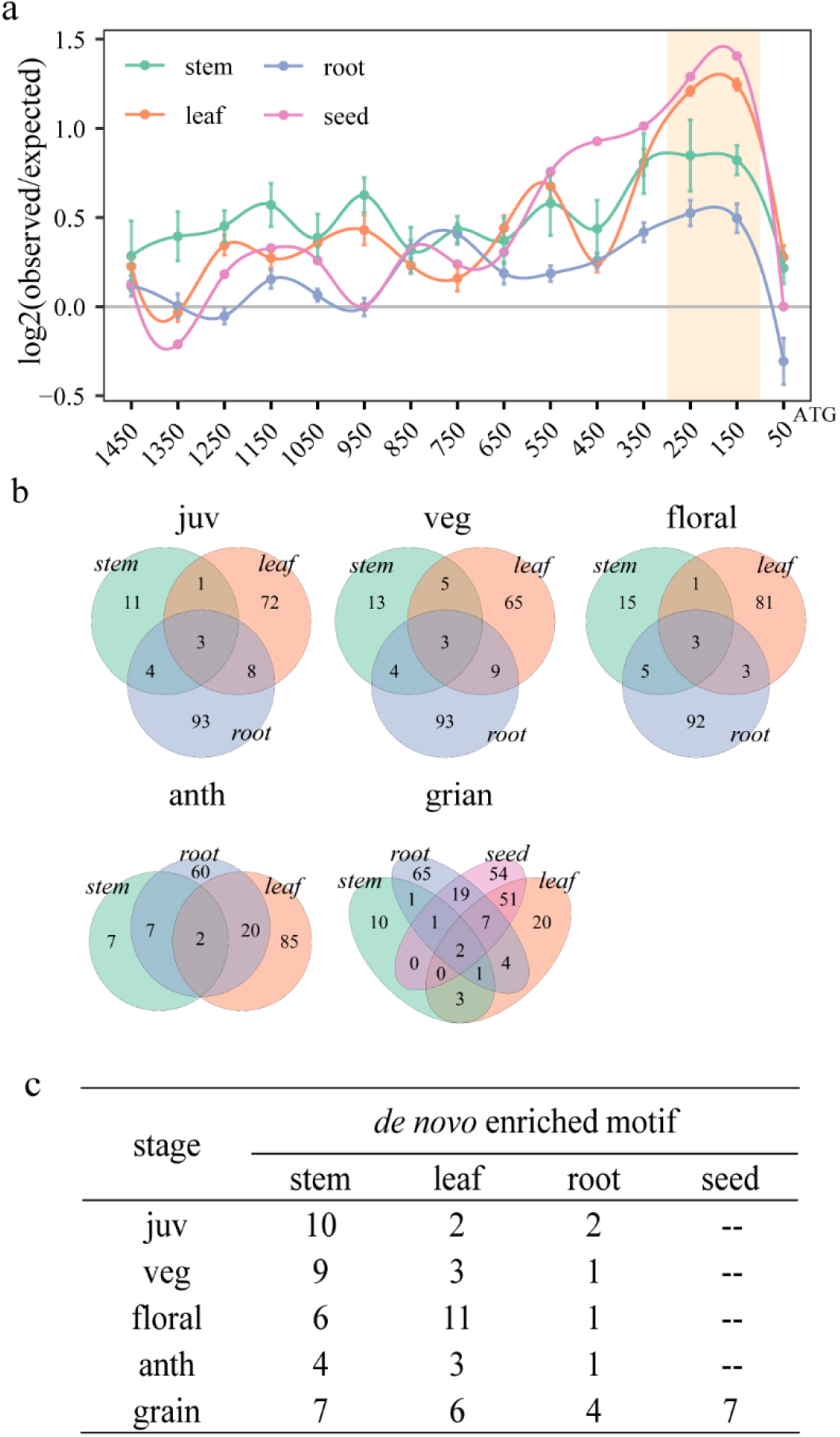
Enriched known and *de novo* motifs in the promoter regions of organ-specific genes. (a) Positional distribution of enriched known motifs along promoter region. The ratio of observed to expected occurrences (observed/expected) reflects motif positional preferences within the promoter region. (b) Venn diagram of unique and shared enriched known motifs identified from organ-specific gene promoters at individual developmental stages. (c) The number of *de novo* enriched motifs discovered by MEME (*E-value* < 0.05).

**Figure 8.**
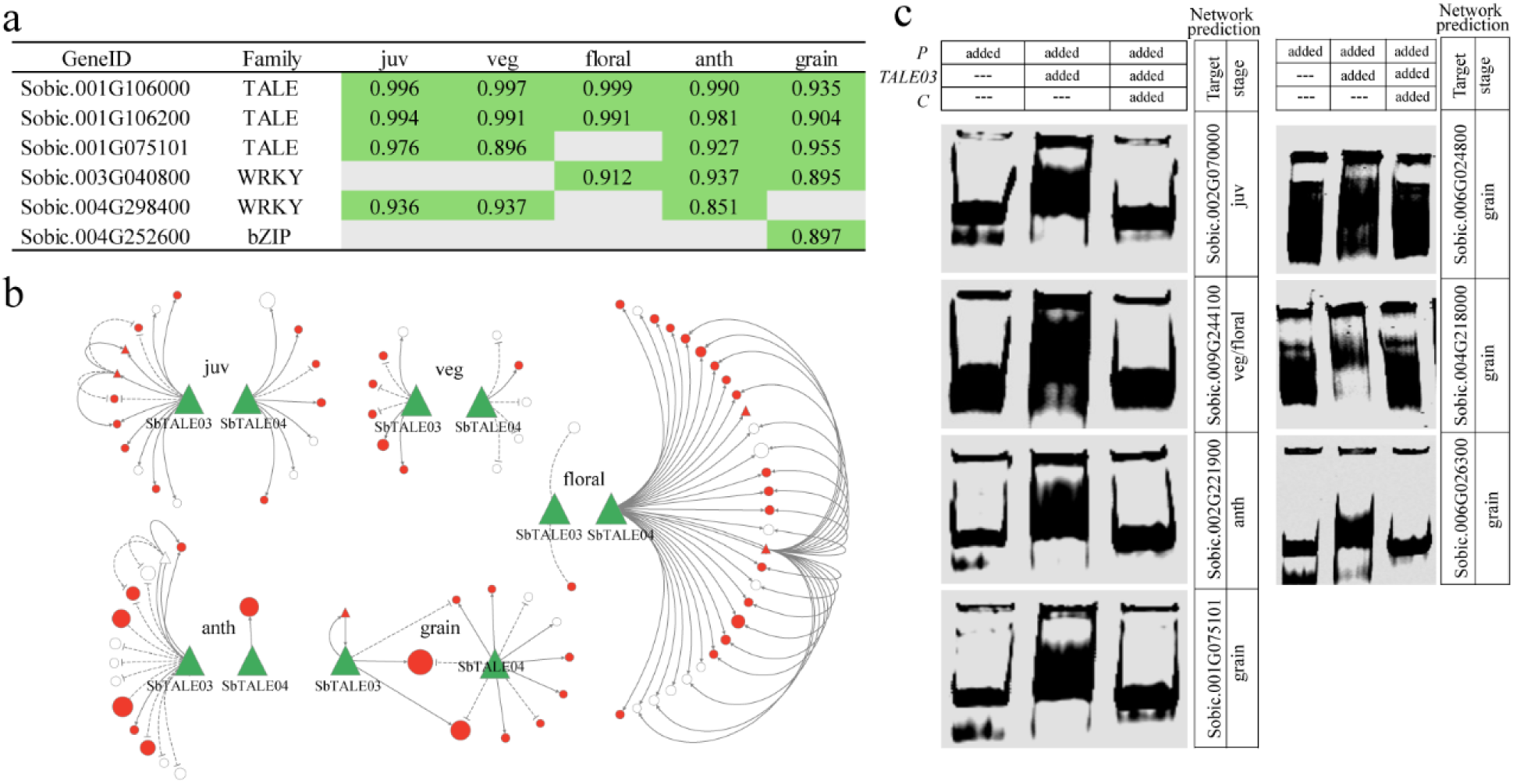
Gene regulatory networks of SbTALE03 and SbTALE04. (a) Evaluation of six ubiquitous stem-specific transcription factors (TFs) for their role as stem hubs across developmental stages. Each TF is classified as a hub (green box) or non-hub (grey box) based on gene significance (GS) values. (b) Topological structure of the regulatory network. Node sizes represent the expression strength of target genes. Red nodes indicate stem-specific genes, while white nodes represent non-stem-specific genes. Solid arrows denote activation (PCC > 0), while dashed lines indicate repression (PCC < 0). Detailed gene identities and network edges are provided in Supplemental Data 10. (c) Electrophoretic mobility shift assay (EMSA) results. *P*: 1.5 kb dsDNA promoter sequence labeled with IRDye 800RS fluorescence (1X concentration); *C*: Unlabeled competitor promoter sequence (100X concentration); *TALE03*: purified protein.

Fewer enriched known motifs were identified in the promoters of stem-specific genes than in those of other organs, and this pattern was consistently observed at all five developmental stages (Figure 7b), possibly due to the limited representation of stem-derived regulatory elements in current motif databases. We next performed *de novo* motif discovery using MEME. For each organ type, only a limited number of *de novo* enriched motifs were identified (Figure 7c). Among those found in the promoters of stem-specific genes, 19 (53%) exhibited low similarity (Pearson correlation coefficient < 0.48) to known motifs (Figure S5). These low-similarity motifs could represent true *de novo* elements that are not currently captured in existing motif databases (Supplemental Dataset 5). Given their novelty and potential functional relevance, these motifs may serve as additional candidates for future studies on sorghum stem-specific gene regulation. The presence of these known and *de novo* enriched motifs in the promoters of all six ubiquitous stem-specific transcription factors underscores their importance in stem-specific gene expression and stem functions (Figure S6).

Comparison of enriched known motifs occurring within organ-specific gene promoters revealed 7–15 motifs per stage of plant development that were uniquely enriched in stem-specific promoters (Figure 7b and Supplemental Dataset 6). Among these motifs, 6–9 of these motifs were enriched in a single stage of development (Figure S7). Only one motif (MA0965.1) was consistently detected in both the juvenile and vegetative stages (before floral initiation), and one motif (MA0937.1) was found in both anthesis and grain maturity stages (after floral initiation). These stem-specific motifs may contribute to defining stem functions that are distinct from those of leaf and root organs, and to modulating stem activity in response to developmental cues.

### Regulatory networks of stem hub transcription factors

Identifying transcription factors (TFs) central to stem functionality is a critical step toward selecting candidates for downstream functional studies and engineering applications. We assessed gene significance (GS) values of TFs within stem-important WGCNA co-expression modules and identified 84 stem hub TFs (GS > 0.8 and *p* < 0.05; Supplemental Dataset 7).

Among these, SbTALE03 and SbTALE04 were the only two TFs consistently classified as stem hubs across all developmental stages (Figure 8a). Their prominent positions within co-expression networks, combined with high stem-specific expression (Figure 4, 5 and 6), underscore their fundamental importance in sorghum stem biology.

To further investigate the regulatory influence of SbTALE03 and SbTALE04, we constructed gene regulatory networks (GRNs) by integrating transcription factor binding site (TFBS) predictions with co-expression strength (PCC) and mapped these interactions within the modules containing SbTALE03 and SbTALE04, which co-occur in the same module at most developmental stages (Supplemental Dataset 7). Although this network construction strategy limited the total number of predicted targets, it identified high-confidence regulatory interactions that offer reliable insights into gene function in sorghum stem organ.

During the juvenile and early adult vegetative stages, stem internode growth is suppressed, allowing production of a stack of nodes from which crown roots emerge. During this phase of development stems increase from 2 mm to 1 cm in size due to the activity of the shoot apical meristem. Analysis of RNA-seq data from stem tissue at this stage of development showed that SbTALE03 and SbTALE04 were predicted to regulate genes associated with meristematic cell maintenance (Supplemental Dataset 8). For instance, SbTALE03 is linked with *Sobic.001G448000*, a homolog of AINTEGUMENTA (ANT), a member of the AINTEGUMENTA-LIKE (AIL) transcription factor family that plays a key role in meristem maintenance and organogenesis. In addition, both SbTALE03 and SbTALE04 were predicted to negatively regulate genes involved in cell wall expansion and lignin biosynthesis, including *Sobic.003G161700* (a homolog of Wall-Associated Kinase 5) and *Sobic.006G011700* (a

Dirigent-like protein), respectively. These observations suggest a shared role of both SbTALE03 and SbTALE04 in preventing secondary cell wall differentiation and promoting continued cell proliferation and growth from a more meristematic, undifferentiated state.

In contrast, GRNs constructed for more mature stem organs, which were collected from floral initiation through grain maturity and excluded the SAM, revealed a functional shift in the regulatory roles of SbTALE03 and SbTALE04. Unlike the early developmental stages where both genes displayed similar regulatory strength, SbTALE03 became more active at anthesis, regulating a greater number of targets than SbTALE04 (Figure 8b), whereas SbTALE04 exhibited higher activity during floral initiation (Figure 8b). Specifically, at floral initiation, SbTALE04 primarily regulated genes involved in cell-cell communication and defense response, including several receptor-like kinases (*Sobic.001G116600*, *Sobic.001G389600*, *Sobic.002G006100*) and a leucine-rich repeat protein (*Sobic.001G021700*) (Supplemental Dataset 8). At anthesis, SbTALE03 was associated with the regulation of hormone biosynthesis (e.g., *Sobic.002G221900*, ETHYLENE-FORMING ENZYME) and membrane lipid signaling (e.g., *Sobic.002G235900*, encoding a phosphoinositide kinase) (Supplemental Dataset 8). At grain maturity, both SbTALE03 and SbTALE04 were implicated in epigenetic gene silencing, with shared regulation of *Sobic.004G218000*, encoding a histone methyltransferase (Supplemental Dataset 8).

These findings suggest that SbTALE03 and SbTALE04 play distinct yet complementary roles in the transcriptional regulation of sorghum stem development, shifting from meristem maintenance in early growth stages to more specialized processes, including signaling and defense as well as epigenetic control during stem maturation.

To experimentally validate GRN predictions, we tested the physical interaction between protein and the promoter regions of several predicted target genes, using electrophoretic mobility shift assays (EMSA). We were only able to purify SbTALE03 protein using the *E. coli* expression system for EMSA analysis (Figure S8). Although SbTALE03 and SbTALE04 share high protein sequence similarity (*E-value* = 2e-93), the presence of unique domains in SbTALE04, specifically, the chloramphenicol acetyltransferase–like domain and the vitellinogen open beta-sheet subdomain, may contribute to poor solubility or aggregation with endogenous *E. coli* proteins hindering efficient elution during purification. Our EMSA results for SbTALE03 showed that the protein bound all seven tested promoter DNA sequences *in vitro* (Figure 8c).

Although we were unable to experimentally test SbTALE04 or assess all predicted targets from GRN, the successful validation of SbTALE03 binding and the consistency of results support the accuracy of our GRN construction.

## Discussion

Genome-wide analysis of transcriptome data from sorghum organs revealed that fewer stem-specific genes were identified compared to other organ types (Figure 1 and S1). The number of stem-specific genes is only 24–62% of leaf-specific genes and 21–64% of root-specific genes, respectively, depending on developmental stage. This trend is consistent with findings by Shakoor et al. [49], who reported fewer conserved stem-specific genes across three sorghum genotypes (forage, sweet, and biomass) compared to conserved leaf- and root-specific genes. One possible explanation lies in the developmental origin and organization of meristems. Angiosperms possess two primary meristems responsible for organogenesis and long-term growth: the shoot apical meristem (SAM), which gives rise to aerial organs including leaves, stems, and flowers [70], and the root apical meristem (RAM), which has evolved to exclusively generate roots [71] and a specialized transcriptome from those of leaves and stems. The SAM is organized into three clonally distinct layers L1, L2, and L3 [72] whose daughter cells differentially impact organ-specific gene expression. The L1 gives rise to the epidermis [72], which consists of a single-cell layer and constitutes only a small proportion of total organ mass. As such, its contribution to bulk transcriptome profiles is minimal. Meanwhile, L2-derived subepidermal cells form the majority of the mesophyll in leaves [72], a highly specialized organ for photosynthesis. The functional and structural specificity of leaf mesophyll may require a broader set of uniquely expressed genes, thereby contributing to the high number of leaf-specific genes. In contrast, the stem organ is primarily derived from the innermost L3 layer, which also gives rise to the vascular bundles found throughout shoot-derived organs [72–74]. Because L3-derived cells contribute broadly to multiple organs, genes expressed in these cells often participate in shared transcriptional programs and therefore would not be classified as strongly stem-specific. The discovery of SbTALE04 (Sobic.001G106200) as one of the small number of genes specifically expressed in sorghum stems supports the above hypothesis. SbTALE04 is homologous to the maize gene *KNOTTED1* (*E-value* of 1.8e-74 with 92% sequence identity), which is primarily expressed in the L3 layer of the shoot apical meristem and functions to maintain L3 identity [75,76]. Thus, it is not surprising that a gene with a potentially important role in specification of the L3 layer is preferentially-expressed in stems.

In addition to SbTALE04, SbTALE03 (Sobic.001G106000), which shares 83% sequence identity with maize *KNOX3* (*E-value* = 2.8e-170), was also experimentally validated as stem-specific in our study (Figure 5 and 6). This expression pattern is consistent across diverse sorghum genotypes, including grain, sweet, and bioenergy types (Figure 3 and 4). Thus, SbTALE03 and SbTALE04 are promising sources of endogenous promoters for driving stem-preferred transgene expression in sorghum. Sorghum is a key feedstock for bioenergy and bioproducts, yet an ideal engineered ideotype remains to be realized [19]. One major challenge in plant engineering is the widespread use of constitutive promoters, which often cause yield penalties or unintended organ effects [77]. In contrast, organ-specific promoters, such as those from SbTALE03 and SbTALE04, offer an opportunity to spatially restrict transgene expression and reduce off-target impacts. Further experimental validation is needed to confirm their developmental regulatory roles.

The binding of transcription factors to their regulatory motifs is a critical step in directing gene expression and establishing distinct expression profiles among organ types [29]. Our computational analysis revealed an enrichment of organ-specific regulatory motifs within the 300 bp region upstream of the ATG (Figure 7a). This positional bias aligns with previous findings in yeast [78], humans [79], and Arabidopsis [69,80,81]. It emphasizes the well-established importance of the core promoter region (approximately TSS ± 40 bp) in the assembly of the transcriptional machinery, but also the potential regulatory significance of the region near the 5′ untranslated region (5′ UTR) in mediating organ-specific gene expression. Studies have demonstrated that sequence variation within the 5′ UTR can influence organ-specific gene expression. For instance, alternative splicing of 5′ UTRs has been shown to regulate gene expression in an organ-specific fashion, and differential use of untranslated regions can lead to variation in gene expression profiles across different organs [82]. These findings underscore the significant regulatory functions of the 5′ UTR in controlling gene expression and its impact on organ-specific expression patterns. One limitation of deriving organ-specific motifs from co-expressed organ-specific genes is that hard clustering methodologies may fail to capture subtle co-expression patterns dictated by unique sets of motifs. Additionally, relying solely on a single motif discovery tool can introduce biases leading to false positive or negative regulatory elements. To address these challenges, previous studies have incorporated more expression datasets encompassing broader biological variations, applied soft clustering approaches, and integrated multiple motif discovery tools to improve the identification of organ- or condition-specific regulatory motifs [69,83,84].

Our constrained and experimentally validated GRN analysis highlights a stage-specific regulatory shift in SbTALE03 and SbTALE04, from maintaining pluripotent, undifferentiated meristematic cell identity during early development (juvenile and vegetative stages) to orchestrating more specialized functions upon the initiation of reproductive growth (after floral initiation). The early-stage regulatory roles of these transcription factors align with the well-characterized functions of KNOTTED1-like homeobox (KNOX) proteins. For example, KN1, the founding member of the KNOX family in maize [75,76], is known to maintain meristem indeterminacy [85,86] by modulating hormone pathways [87,88]. BREVIPEDICELLUS (BP; AT4G08150) in Arabidopsis, homologous to both SbTALE03 and SbTALE04, similarly promotes shoot apical meristem indeterminacy and negatively regulates lignin biosynthesis and secondary cell wall thickening [89]. The observation that sorghum SbTALE03 and SbTALE04 regulate genes functionally annotated for meristem maintenance and delayed cell wall maturation provides evidence for the evolutionary conservation of homeodomain protein functions across plant species. In line with their roles in hormone-related transcriptional control, SbTALE04 is also predicted to regulate *Sobic.004G329000* during the juvenile stage, a gene encoding BRS1, which functions in the brassinosteroid signaling pathway [90].

Notably, SbTALE03 and SbTALE04 in sorghum, and their maize homologs KNOX3 and KN1, respectively, are arranged as tandem duplicates. The presence of these gene pairs in both species suggests that the duplication event occurred prior to their divergence. Following floral initiation, the regulatory networks indicate that SbTALE03 and SbTALE04 in sorghum diverge toward more specific, developmentally responsive roles in mature stem organ. Similarly, in maize, KN1 and KNOX3 function in distinct regulatory networks and KN1 loss-of-function does not affect KNOX3 expression or activity [87]. These parallels suggest that regulatory divergence of this tandem gene pair likely began soon after duplication and has been maintained in both species, consistent with subfunctionalization as a mechanism contributing to the retention of both genes.

Although our GRN construction method integrated multiple layers of biologically relevant information, including transcription factor binding site (TFBS) predictions and co-expression relationships, it still has important limitations. In particular, it lacks regulatory data that capture chromatin context and transcription factor (TF) occupancy, which are known to be both species- and organ-specific [29,91–93]. The TF-binding information used here was primarily derived from Arabidopsis [94] or inferred based on protein DNA binding domain similarity [95], which may not fully reflect the regulatory landscape in sorghum due to evolutionary divergence. Although methods such as DAP-seq, ChIP-seq, and ATAC-seq can provide valuable and complementary insights into the regulatory architecture, such datasets are currently unavailable for sorghum. The future availability of sorghum-specific regulatory data would significantly improve the accuracy and biological relevance of inferred GRNs.

## Conclusion

This study presents a comprehensive, stage-resolved transcriptomic and regulatory analysis of *Sorghum bicolor*, with a particular focus on stem-specific gene expression, function, and regulation. By integrating RNA-seq re-analysis, co-expression networks, gene regulatory network construction, motif discovery, and empirical validation, we identified a core set of organ-specific genes with ubiquitous stem specificity across genotypes and developmental stages.

Among these, two homeodomain transcription factors, *SbTALE03* and *SbTALE04*, were experimentally validated and shown to play key roles in stem developmental regulation. Our findings provide new information in transcriptional and cis-regulatory architecture underlying sorghum stem function and development. The identified stem-specific genes, motifs, and regulatory interactions serve as resources for future studies and offer promising targets for precision engineering aimed at improving stem biomass composition and productivity in bioenergy sorghum.

## Declarations

### Ethics approval and consent to participate

Not applicable

### Consent for publication

All authors have read and approved the final manuscript and consent to its publication.

### Availability of data and materials

All RNA-seq datasets analyzed in this study are publicly available from the NCBI Sequence Read Archive (accessions SRA558272, SRA558514, and SRA558539) and the JGI Genome Portal (identifiers 1258721 and 1258409). All supporting files are available in the supplemental materials.

### Competing interests

The authors declare that they have no competing interests.

## Funding

This work was funded by the DOE Center for Advanced Bioenergy and Bioproducts Innovation (U.S. Department of Energy, Office of Science, Biological and Environmental Research Program under Award Number DE-SC0018420). Any opinions, findings, and conclusions or recommendations expressed in this publication are those of the author(s) and do not necessarily reflect the views of the U.S. Department of Energy.

## Authors’ contributions

JF, MHA, AMC, KS, and JEM conceived the project and designed the experiments. AMC and KS secured funding. JEM and SPM contributed experimental data. JF performed all major data analyses. BJ conducted the RNAscope validation, and YL carried out the EMSA experiments. BM investigated the gene expression patterns of *SbTALE03* and *SbTALE04*. EK performed the phylogenetic analysis. KB generated RNA-seq data from stem and leaf organs across multiple genotypes. JF, BJ, YL, EK , SPM, JEM, KS, and AMC wrote the manuscript. All authors reviewed and approved the final manuscript.

## Acknowledgements

Not applicable

---

**Figure S1.**
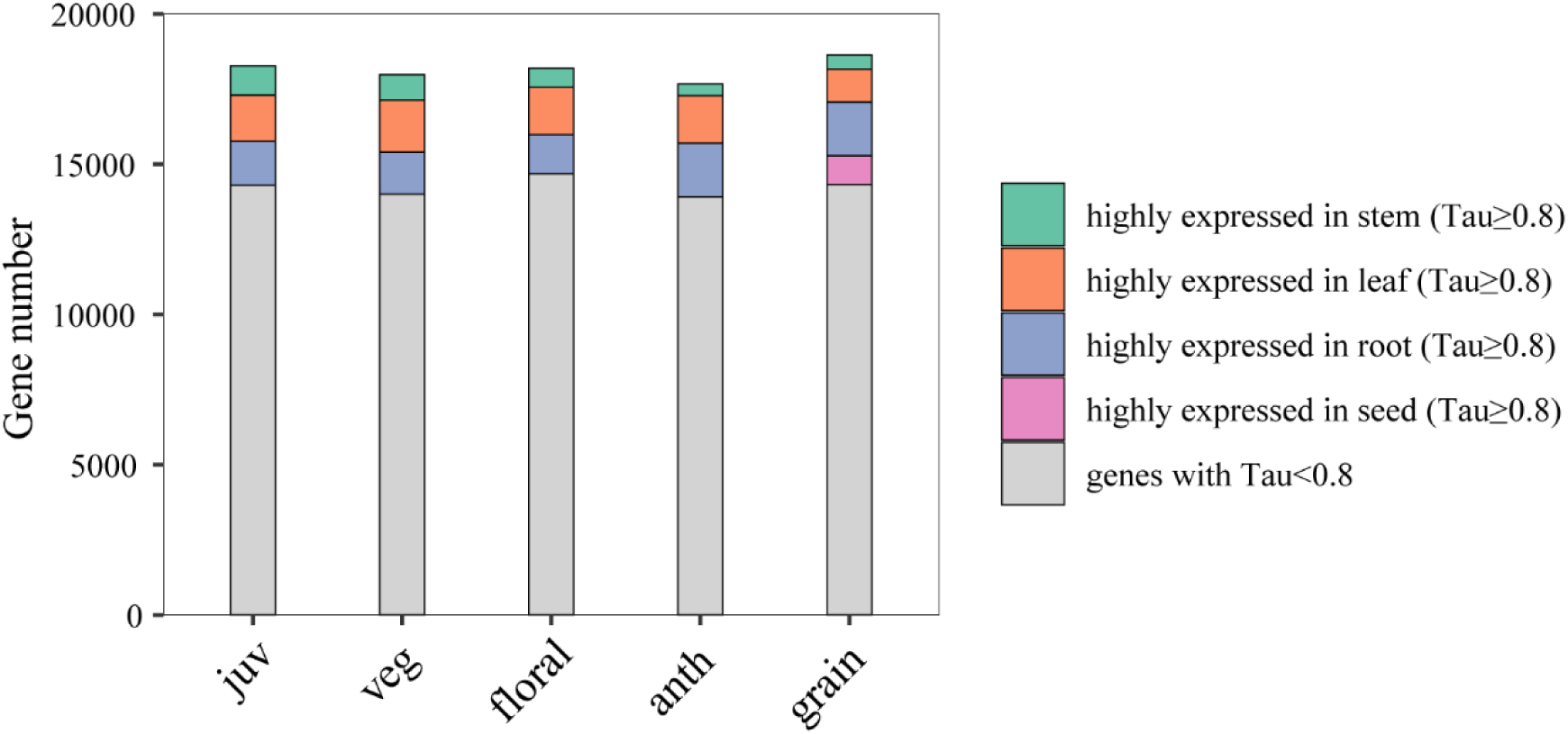
Genes highly expressed in specific organ types based on Tau metric.

**Figure S2.**
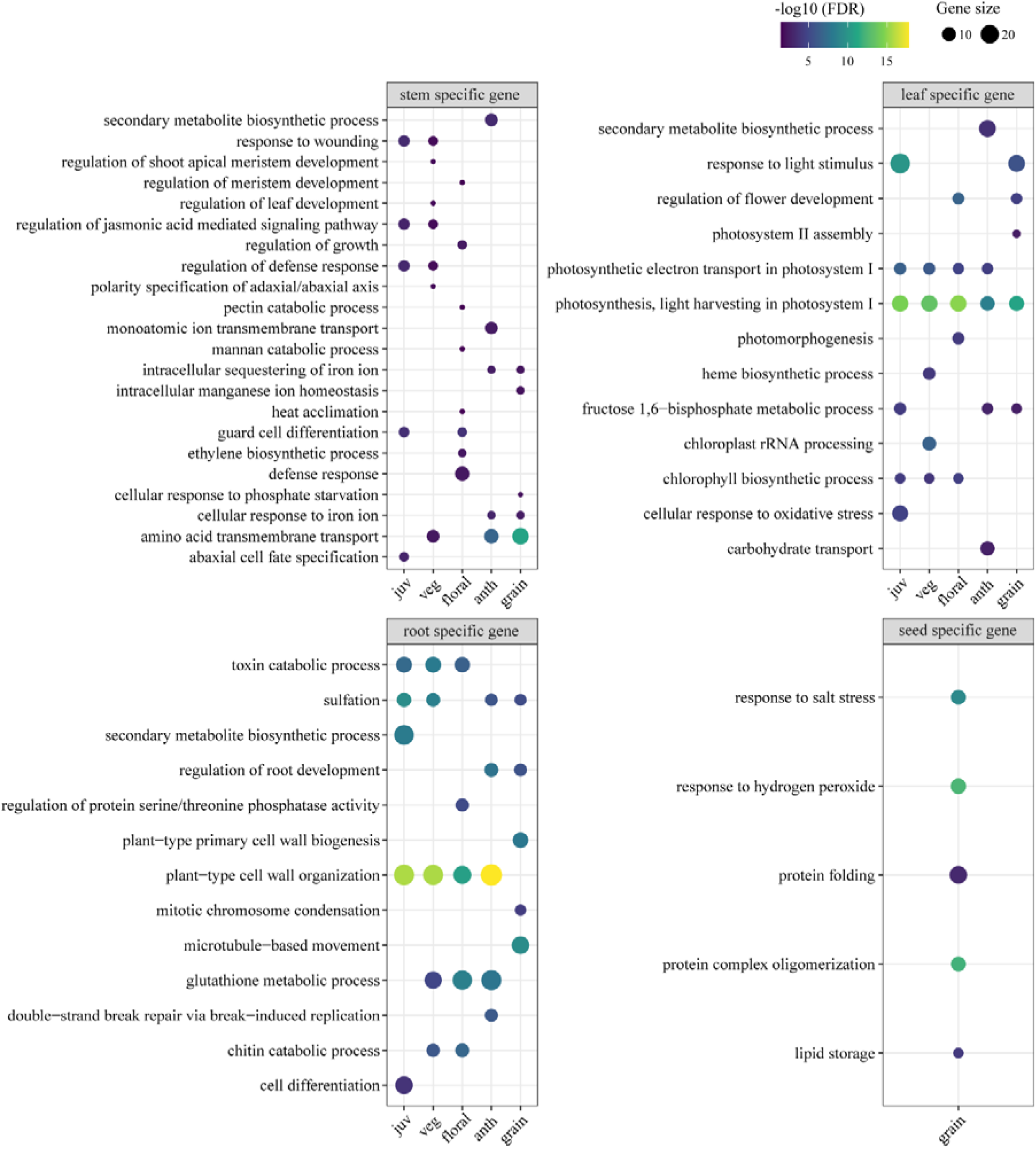
Enriched biological functions of organ-specific genes. The top five most specific enriched Gene Ontology (GO) Biological Process (BP) terms for each developmental stage are shown, as determined by the method described in the Methods section. Complete enrichment results are provided in Supplemental Dataset 3.

**Figure S3.**
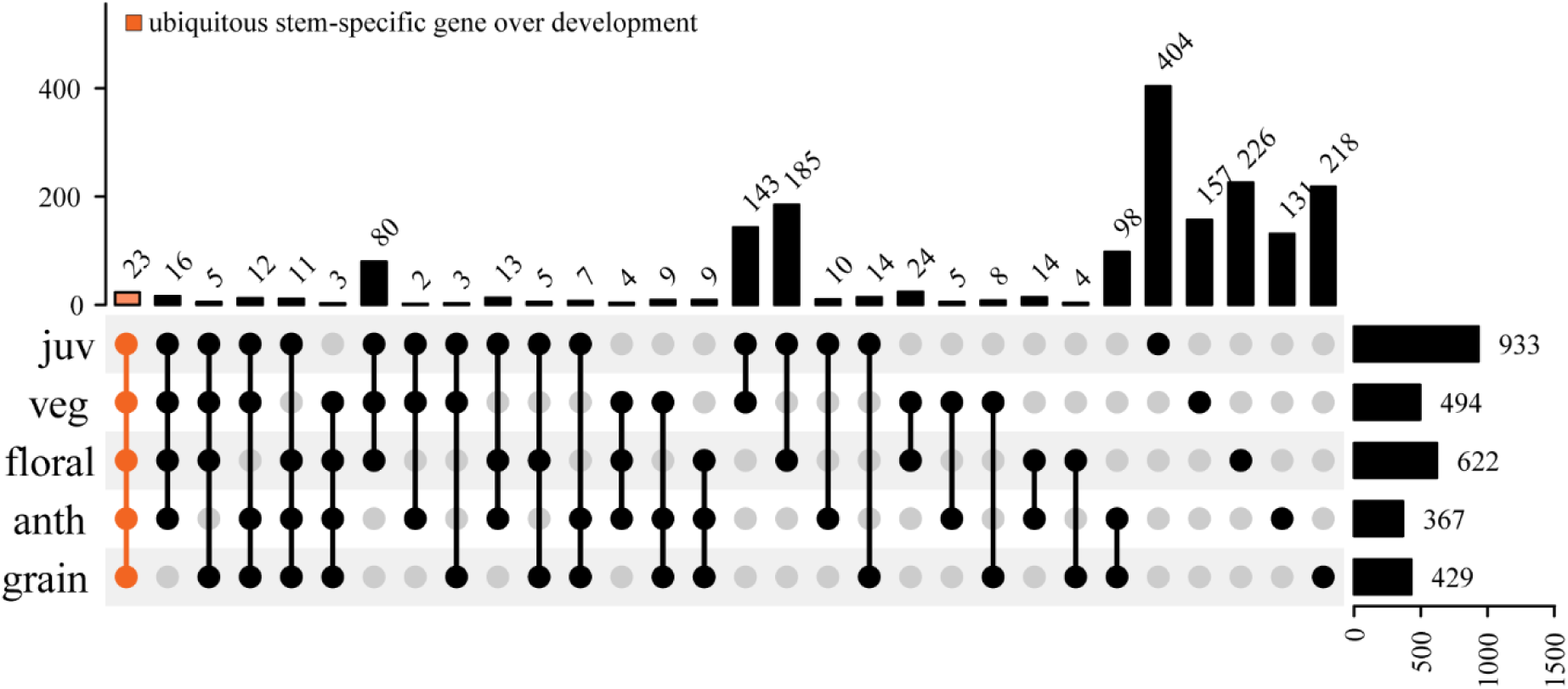
Unique and common stem-specific genes across developmental stages.

**Figure S4.**
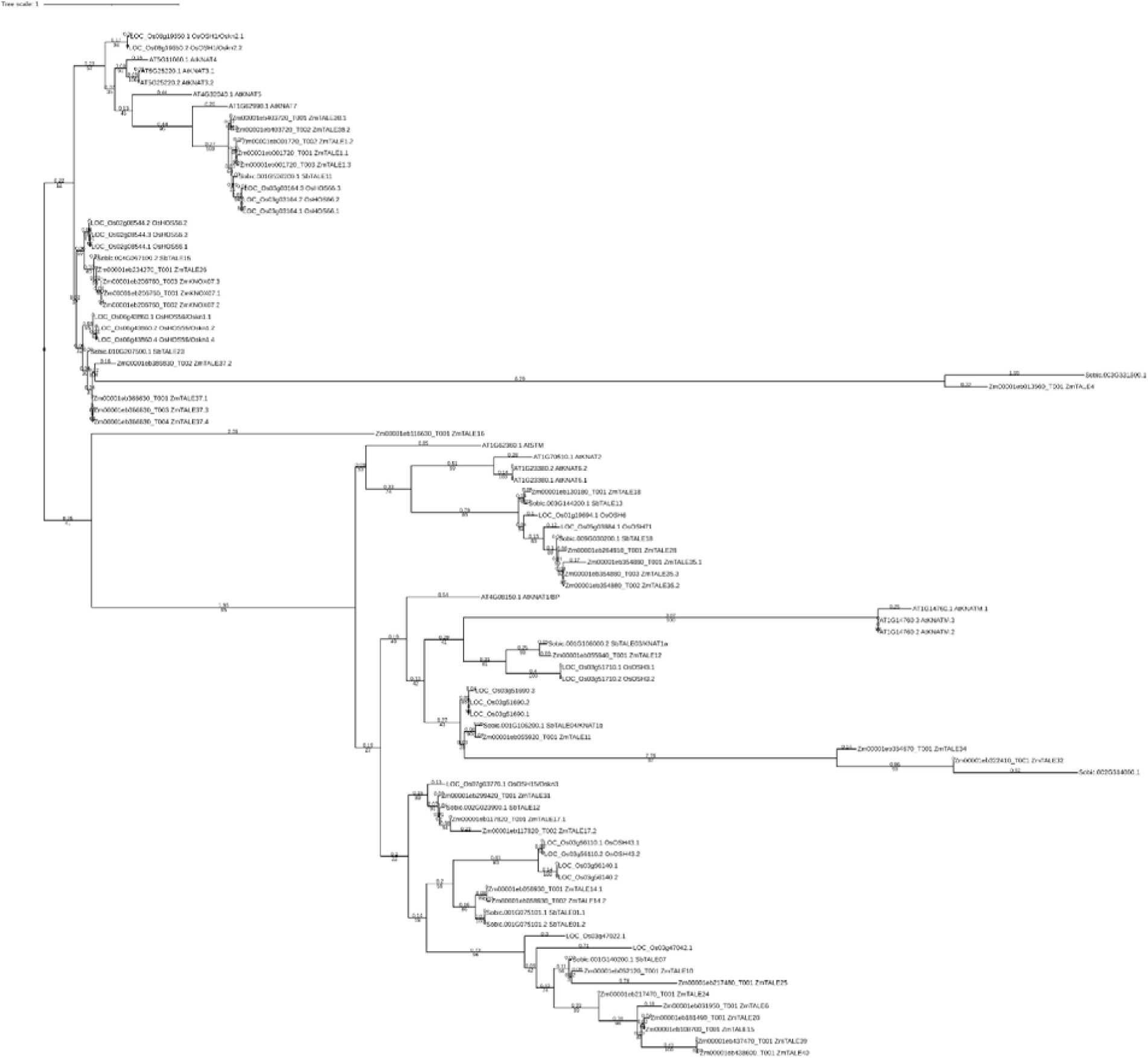
Phylogenetic tree of KNOX (knotted-like homeobox) family proteins from Sorghum, Rice, Maize, and Arabidopsis. The phylogenetic tree was generated with 1,000 bootstrap replicates and midpoint-rooted for visualization.

**Figure S5.**
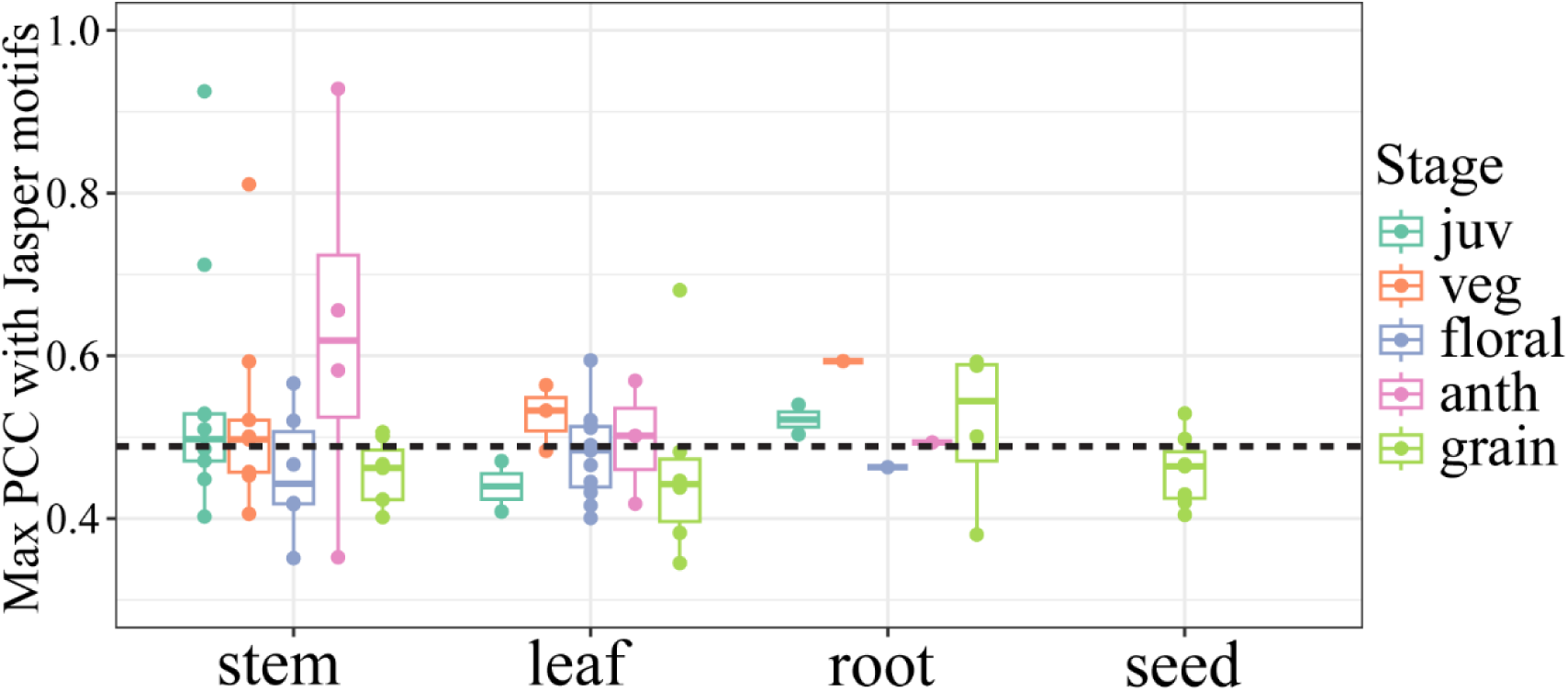
Maximum Pearson correlation coefficient (PCC) between *de novo* enriched motifs and known motifs. The red dashed line (PCC = 0.49) represents the 95th percentile of PCCs in the known motif database, serving as a cutoff for motif similarity. PCC values less than 0.49 indicate distinct motifs that are not represented in current motif database.

**Figure S6.**
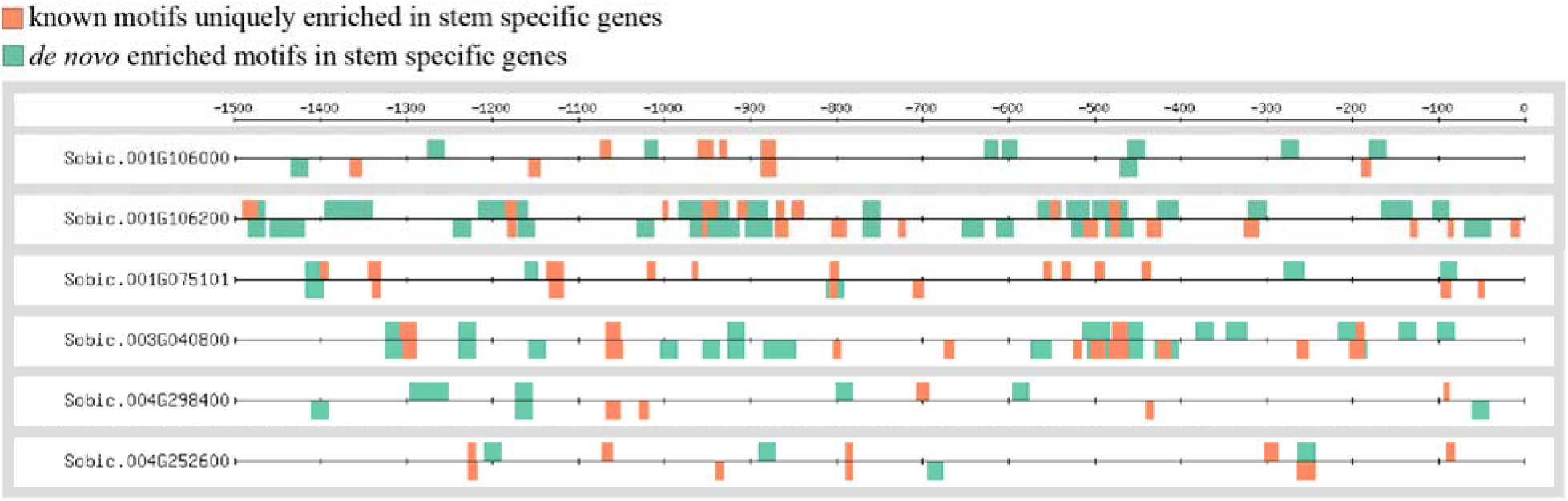
Promoter motif distribution of six ubiquitous stem-specific transcription factors. Motif identities are provided in Supplemental Dataset 6 (known motifs uniquely enriched in stem-specific gene promoters) and Supplemental Dataset 5 (*de novo* enriched motifs in stem-specific gene promoters).

**Figure S7.**
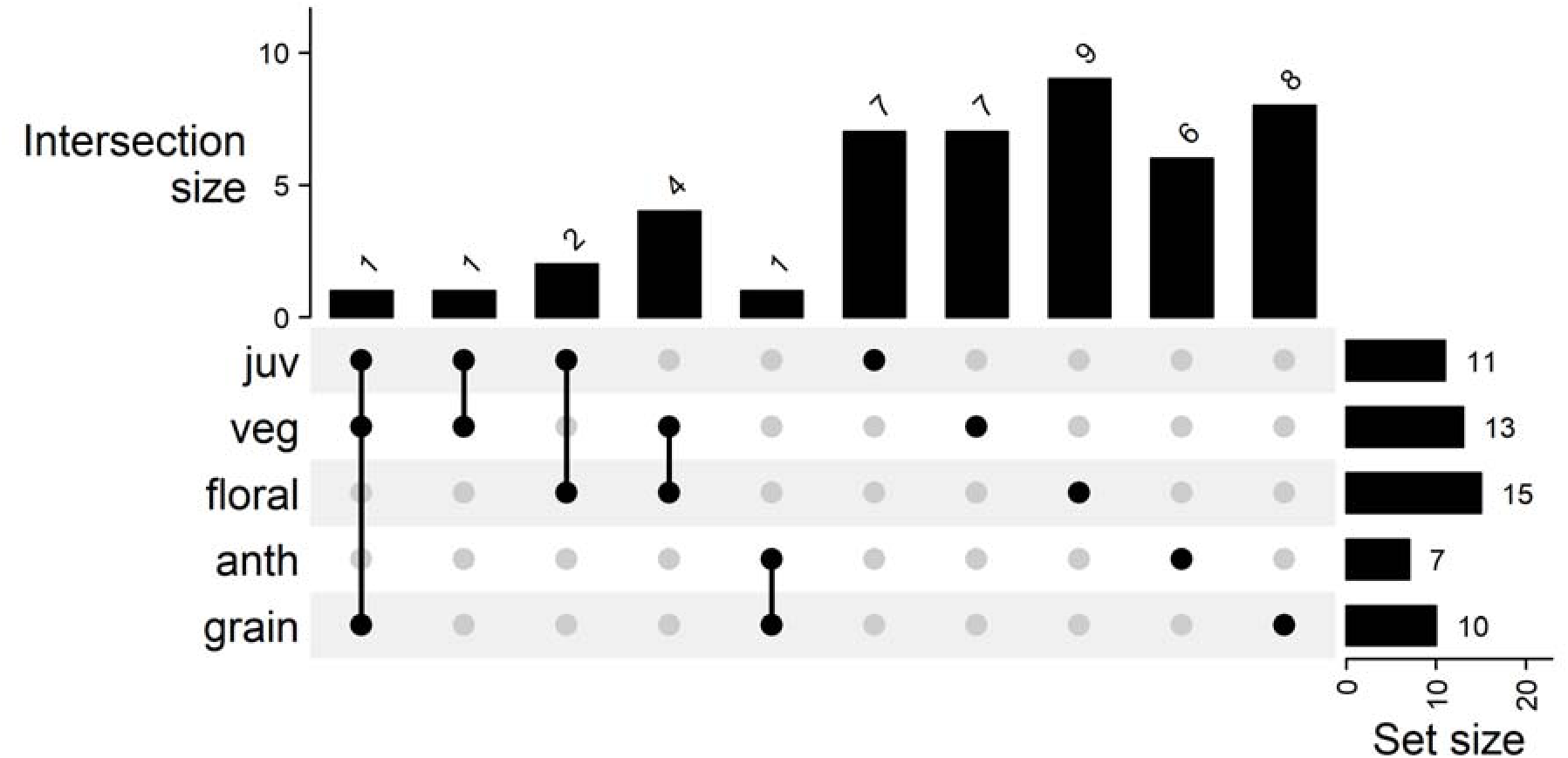
UpSet of uniquely enriched known motifs in stem specific gene promoter region compared to leaf and root. The plot illustrates stage-dependent enrichment of these motifs in the promoter regions of stem-specific genes.

**Figure S8.**
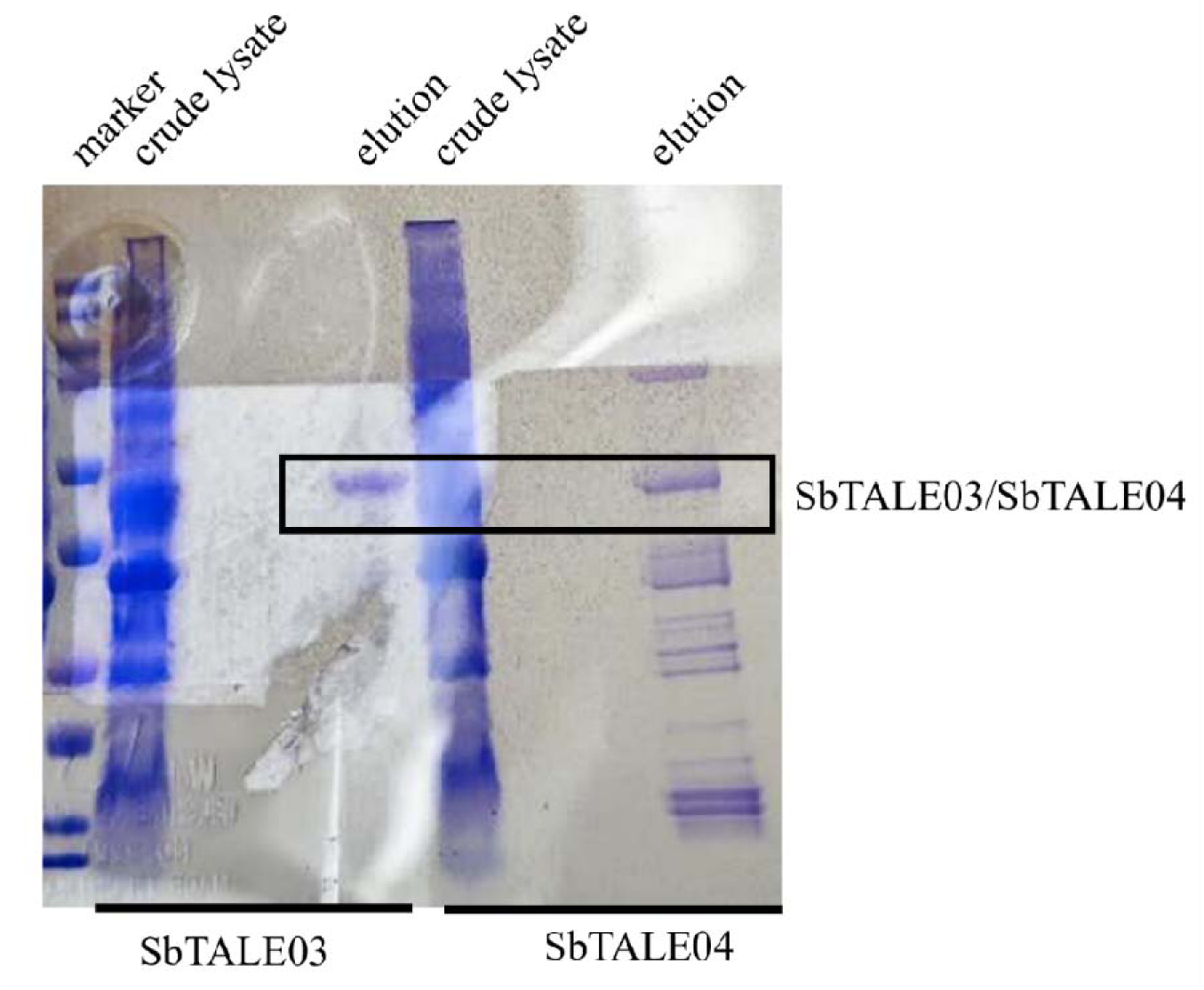
Co-purification of contaminating proteins during SbTALE04 expression in the *E. coli* system.

**Figure S9.**
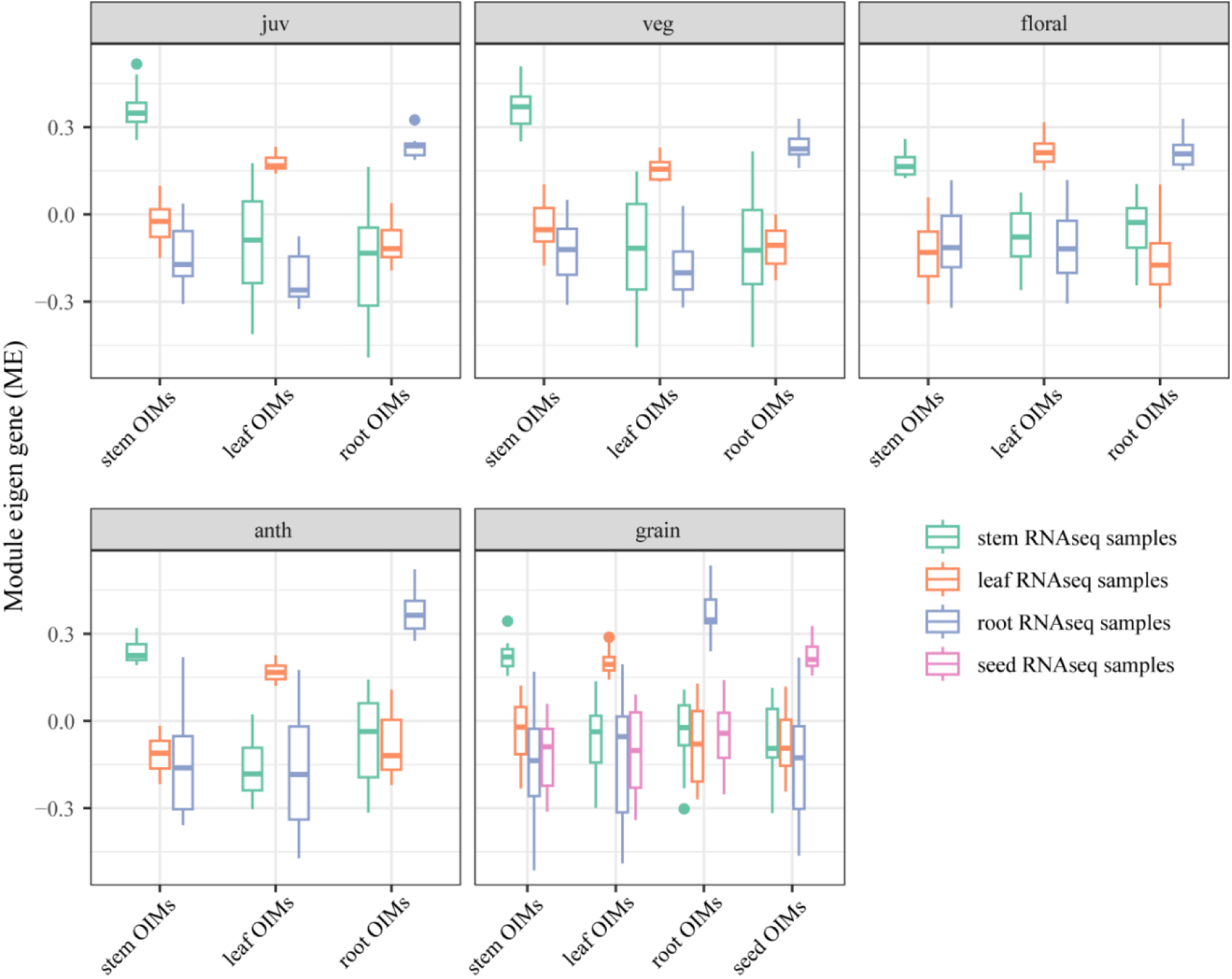
Organ-important modules (OIMs) showed high module eigengene values in corresponding tissue type samples.

**Figure S10.**
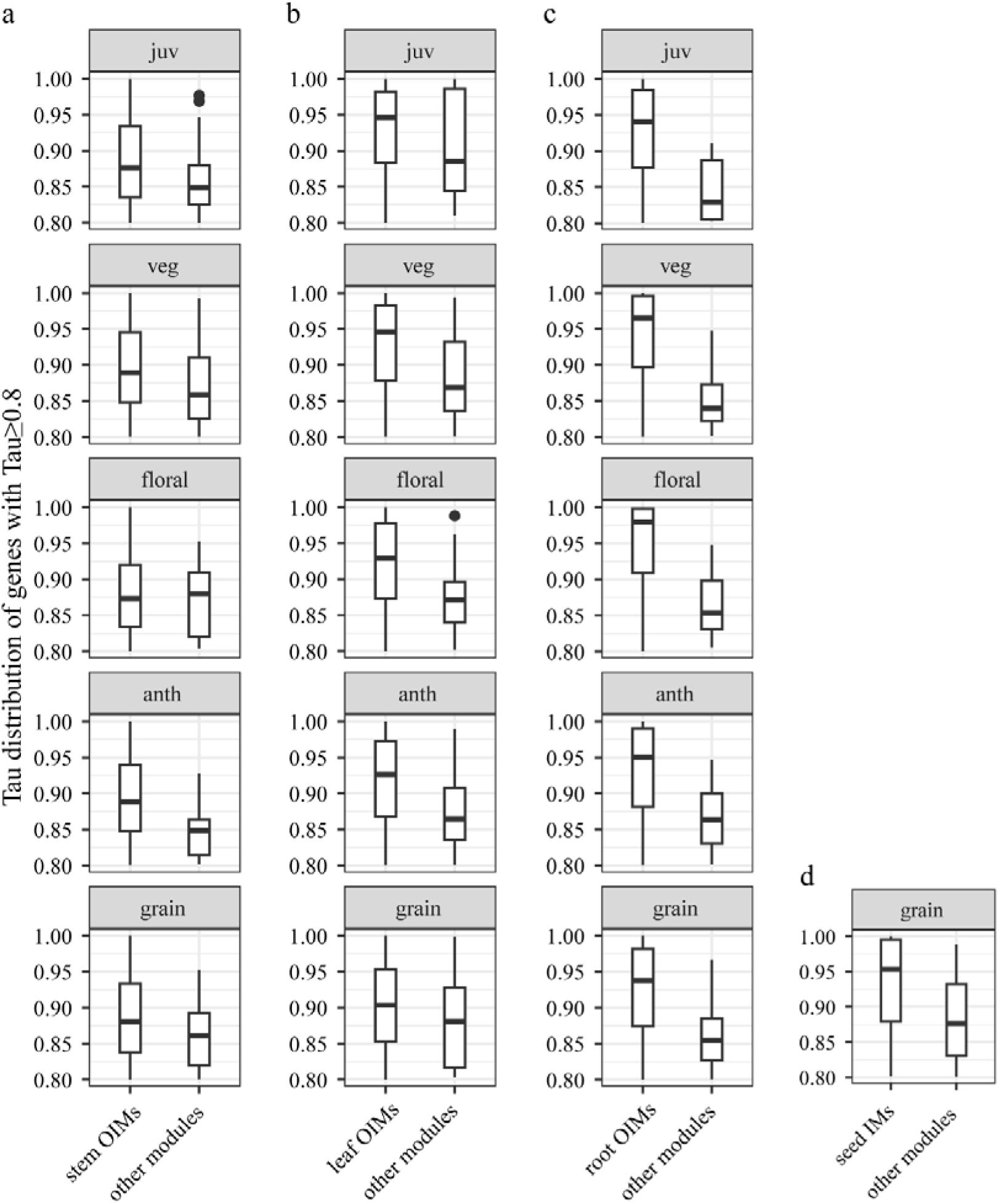
Tau value comparison between organ-important modules (OIMs) and other modules. Genes involved in this comparison are organ-specific genes derived from Tau metric (≥ 0.8). (a– d) are stem (a), leaf (b), root (c), and seed (d).

## Supplemental Datasets

**Supplemental Dataset 1.** BTx623 RNA-seq samples used in Tau analysis and WGCNA

**Supplemental Dataset 2.** Organ-specific genes identified by Tau analysis and WGCNA

**Supplemental Dataset 3.** Enriched GO and KEGG for organ-specific genes

**Supplemental Dataset 4.** Stem-specific genes over development

**Supplemental Dataset 5.** MEME de novo enriched motifs in stem-specific gene promoters

**Supplemental Dataset 6.** Enriched known motifs in organ-specific gene promoters

**Supplemental Dataset 7.** WGCNA hub transcription factors for organ identity##

**Supplemental Dataset 8.** SbTALE03 and SbTALE04 gene regulatory networks##

**Supplemental Dataset 9.** WGCNA module identity and gene significance

**Supplemental Dataset 10.** WGCNA module importance for organ identity

## Notes

### Competing Interest Statement

The authors have declared no competing interest.

